# Arctic phytoplankton spring bloom diversity across the marginal ice zone in Baffin Bay

**DOI:** 10.1101/2022.03.14.484350

**Authors:** Catherine Gérikas Ribeiro, Adriana Lopes dos Santos, Nicole Trefault, Dominique Marie, Connie Lovejoy, Daniel Vaulot

## Abstract

Phytoplankton under-ice blooms have been recently recognized as an important Arctic phenomenon for global primary production and biogeochemical cycling. Drastic sea-ice decline in both extension and thickness enables the development of early blooms, sometimes hundreds of kilometers beneath the pack ice. Baffin Bay is a semi-enclosed sea where Arctic and North Atlantic water masses interact. It is totally covered by sea-ice by March and ice-free by August/September. In the present work, we investigated the phytoplankton community structure across the marginal ice zone between the ice-free, Atlantic-influenced, east and the ice-covered, Arctic-influenced, west Baffin Bay using 18S rRNA high-throughput amplicon sequencing, flow cytometry cell counting and numerous environmental and biological data collected and compiled in the scope of the Green Edge project. Sampling was performed during June-July 2016 in a total of 16 stations with around 6 depths each. Stations were clustered into “Under Ice” (UI), “Marginal Ice Zone” (MIZ) and “Open Water” (OW) on the basis of its sea ice cover upon sampling. Phytoplankton community structure was analyzed by 18S rRNA metabarcoding with the microdiversity approach. The UI sector was characterized by a shallow nitracline, high pico-phytoplankton abundance and a shared dominance between *Micromonas* and *Phaeocystis* in the 0.2-3 *µ*m size fraction, as well as an increased contribution of Cryptophyceae and non-diatom Ochrophyta in the 3-20 *µ*m size fraction. Several amplicon sequence variants (ASVs) were flagged as indicator for the UI+MIZ sector group, including known ice-associated taxa such as the diatoms *Melosira arctica* and *Pseudo-nitzschia seriata*, but also specific ASVs assigned to the green alga *Micromonas polaris* and the cryptophyte *Falcomonas daucoides*, the silicoflagellate *Dictyocha speculum*, one member of the uncultivated MOCH-2 group, and a *Pterosperma* sp. (green algae) rarely seen in other metabarcoding datasets, including from the Arctic. The OW sector harbored a community adapted to a nutrient-depleted/high light environment, with a significant contribution of the haptophyte*Phaeocystis pouchetii* and big centric diatoms, including several *Thalassiosira* species.

## Introduction

The recognition of the occurrence of an annual under-ice bloom in the Arctic Ocean (Arrigo et al. 2012, 2014) represented a paradigm shift that has impacted the estimates of primary production (Kinney et al. 2020), as well as the understanding of the biogeochemical cycling in the region (Ardyna et al. 2020). The Arctic is undergoing drastic changes directly linked to sea-ice decline in both extension and thickness (Meredith et al. 2019; Serreze et al. 2007), enabling the early development of extensive under-ice blooms (Horvat et al. 2017). A recent model indicates that the photosynthetically active radiation (PAR) transmission trough first and second-year sea ice can now sustain net phytoplankton growth over most of the Arctic by July (Ardyna et al. 2020). The periods when sea-ice is present have been shortened by earlier melting and delayed freezing seasons (Tedesco et al. 2019), impacting also the timing of the characteristic ice-edge phytoplankton spring blooms (Janout et al. 2016; Perrette et al. 2011; Renaut et al. 2018), with cascading effects to higher trophic levels and nutrient fluxes (Leu et al. 2011; Post et al. 2013).

Autotrophic communities from high latitude environments are subjected to a light regime mainly dictated by the seasonally restricted input of solar energy, but also by the sea-ice extension/thickness and snowfall rates, which have a light-attenuating effect (Leu et al. 2015). The sea-ice also provides a complex habitat for the sympagic community (Niemi et al. 2011; Olsen et al. 2017), as well as seeding organisms to the water column during melting (Hardge et al. 2017). The presence of sea-ice and its associated community has been linked to higher abundance and better nutrition of pelagic organisms from higher trophic levels (Hop et al. 2011; Schmidt et al. 2018). The Arctic sea-ice harbors complex communities with many metabolic strategies, where different types of ice present different community structures (Comeau et al. 2013), and may act as a flagellate cyst repository, for example for dinoflagellates such as *Polarella glacialis* (Kauko et al. 2018). Sympagic assemblages have a great potential for the discovery of novel protist taxa (Hardge et al. 2017; Ribeiro et al. 2020). They are now facing an imminent threat due to rapid decline on ice extension: a drastic decrease in sympagic protist diversity has been reported in the Arctic due to the loss of multiyear sea ice, which harbors almost 40% more species than first-year ice (Hop et al. 2020). It is expected that Arctic increasing temperatures, enhanced water column stratification and ocean acidification will also favor specific pelagic populations, such as the pico-sized green alga *M. polaris* (Benner et al. 2019; Hoppe et al. 2018; Li et al. 2009).

Diatoms tend to dominate Arctic sympagic communities and under-ice blooms, especially pennate diatoms from the genera *Nitzschia*, *Fragilariopsis*, *Navicula* and *Cylindrotheca* (Ardyna et al. 2020; Hop et al. 2020; Leu et al. 2015), with also the dominance of *Nitzschia frigida* during polar winter (Niemi et al. 2011). As the snow melts during spring/summer, the formation of melt ponds creates a new habitat which might be connected with the water column below. Melt pond communities are often dominated by flagellates (Mundy et al. 2011), and mixo/heterotrophic groups including Chrysophyceae, Filosa-Thecofilosea, and ciliates (Xu et al. 2020). Bottom-ice communities are rich, characterized by the presence of pennate diatoms and the strand-forming centric diatom *Melosira arctica* (Poulin et al. 2014). The seasonally retreating marginal ice zone is followed by massive phytoplankton blooms developing close to and below the ice edge (Perrette et al. 2011). The pelagic phytoplankton harbors a different diatom community from that of sea-ice (Oziel et al. 2019), with stronger presence of centric diatoms such as *Thalassiosira* and *Chaetoceros*, which are more adapted to lower nutrient concentrations and higher luminosity within the ice-free euphotic zone (Kvernvik et al. 2020; Morando and Capone 2018).

Apart from Bacillariophyta, other groups also play pivotal roles in the Arctic ecosystem. The Arctic pico-phytoplankton (0.2-2 *µ*m) is dominated by the Mamiellophyceae *M. polaris*, *Bathycoccus prasinos* and *Mantoniella* spp. (Joli et al. 2017; Lovejoy et al. 2007; Not et al. 2005). *M. polaris* is often the most abundant (Balzano et al. 2012; Lovejoy and Potvin 2011) and is considered an Arctic sentinel species (Freyria et al. 2021) due to the close relationship of its distribution patterns with temperature (Demory et al. 2019). *Phaeocystis* is a globally distributed haptophyte genus, with a great impact on the carbon and sulfur exchange on the ocean/atmosphere interface (Schoemann et al. 2005). The bloom-forming *P. pouchetii* has a pan-Arctic distribution (Lasternas and Agustı 2010), with blooms detected even under thick snow-covered pack ice (Assmy et al. 2017), and recently reported also in Antarctic waters (Trefault et al. 2021).

Among other perils to the Arctic ecosystem, the “atlantification” phenomena was first reported more than a decade ago (Hegseth and Sundfjord 2008), with hydrographic impacts on water column stratification and sea-ice decline due to increased heat fluxes from Atlantic Water (Polyakov et al. 2017), as well as biological impacts, via intrusion by advection of species of temperate origin (Neukermans et al. 2018; Oziel et al. 2020). Several studies also report a phytoplankton downsizing trend in warmer ocean waters (Hilligsøe et al. 2011; Morán et al. 2010). For example, warm anomalies in the Atlantic Water inflow to the Arctic Ocean seems to shift plankton dominance from diatoms cells to small coccolithophores (Lalande et al. 2013; Smyth et al. 2004).

Baffin Bay is a seasonally ice-covered sea within the Canadian Arctic, with a complex interplay of the Atlantic and Arctic, Pacific-originated water masses (see Material & Methods section). The longitudinal physico-chemical gradient created by this system of water masses results in distinct stratification patterns (Randelhoff et al. 2019) and differential sea-ice melting rates (Tang et al. 2004), greatly impacting food web structure and carbon export (Saint-Béat et al. 2020), as well as the permeability of the sea-ice, influencing brine connectivity and nutrient availability to sympagic algae (Tedesco et al. 2019). Baffin Bay is especially susceptible to drastic environmental changes, with a reported increase of 20 days of the length of the melting season compared to four decades ago (Stroeve et al. 2014), due to ongoing changes such as warming on its eastern subsurface boundaries caused by Atlantic inflow, and freshening trends on its Arctic-influenced sectors (Zweng and Münchow 2006).

In the present work, we used high-throughput amplicon sequencing and a microdiversity approach to investigate how the plankton community structure changes across the marginal ice zone between the Atlantic-influenced east and Arctic-influenced west Baffin Bay. The present study provides important insights on the impact of sea-ice loss on ice-associated pelagic plankton.

## Material and Methods

### Study area

Baffin Bay is a seasonally ice-covered sea within the Canadian Arctic, delimited by Greenland in the west and by Baffin Island in the east, where complex interactions between North Atlantic and Arctic water masses take place. The temperate and salty West Greenland Current (WGC), product of the interaction of North Atlantic waters with the Irminger current, flows northward on eastern Baffin Bay along the Greenland coast, coming through the Davis Strait (Tang et al. 2004). Due to its higher density, the WGC cannot pass through the Canadian Archipelago and recirculates counter-clockwise, interacting with the colder, less-saline, Pacific-originated Arctic waters, flowing southward as the Baffin Island Current (BIC) (Jones et al. 2003; Münchow et al. 2015). Sea-ice formation starts in Baffin Bay during October and covers almost the totality of its area by March, followed by the melting season onset in April, as sea-ice retreats westward until it reaches a minimum extent by August/September (Tang et al. 2004). In the western Baffin Bay, the onset of snow cover melt modulates the termination of the sea-ice algal bloom and the beginning of the under-ice phytoplankton spring bloom, reaching similar magnitudes to its offshore counterparts (Oziel et al. 2019).

### Sampling & DNA extraction

Samples were collected onboard the research icebreaker CCGS *Amundsen* on four longitudinal transects between 68.4*^◦^*N-70*^◦^*N and 56.8*^◦^*W-62.4*^◦^*W, from 9 June to the 2nd July 2016, for a total of 16 sampling stations (Figure 1). Sea water was sampled at six depths within the euphotic layer at each station, using 12-L Niskin bottles attached to a rosette equipped with a Seabird SBE-911plus CTD unit (Sea-Bird Electronics, Bellevue, WA, USA). The list of the sensors attached to the rosette carousel can be found in Bruyant et al. (2022). Three liters of water from each sampling point were pre-filtered with a 100 *µ*m mesh and subsequently filtered with a peristaltic pump through the following sets of polycarbonate filters: 20 *µ*m (47 mm), 3 *µ*m (47 mm), and 0.22 *µ*m (Sterivex™ filters). Filters were placed in cryotubes (except for the Sterivex™), preserved with 1.8 mL of RNAlater™ and stored at −80*^◦^*C until processing. DNA was extracted using ZR Fungal/Bacterial DNA MiniPrep (Zymo Research, Irvine, CA, USA) following instructions from the manufacturer, and its final concentration was measured using PicoGreen™ (Thermo Fisher Scientific, Waltham, MA, USA) with a LabChip GX (Perkin-Elmer, Waltham, MA, USA).

**Figure 1:**
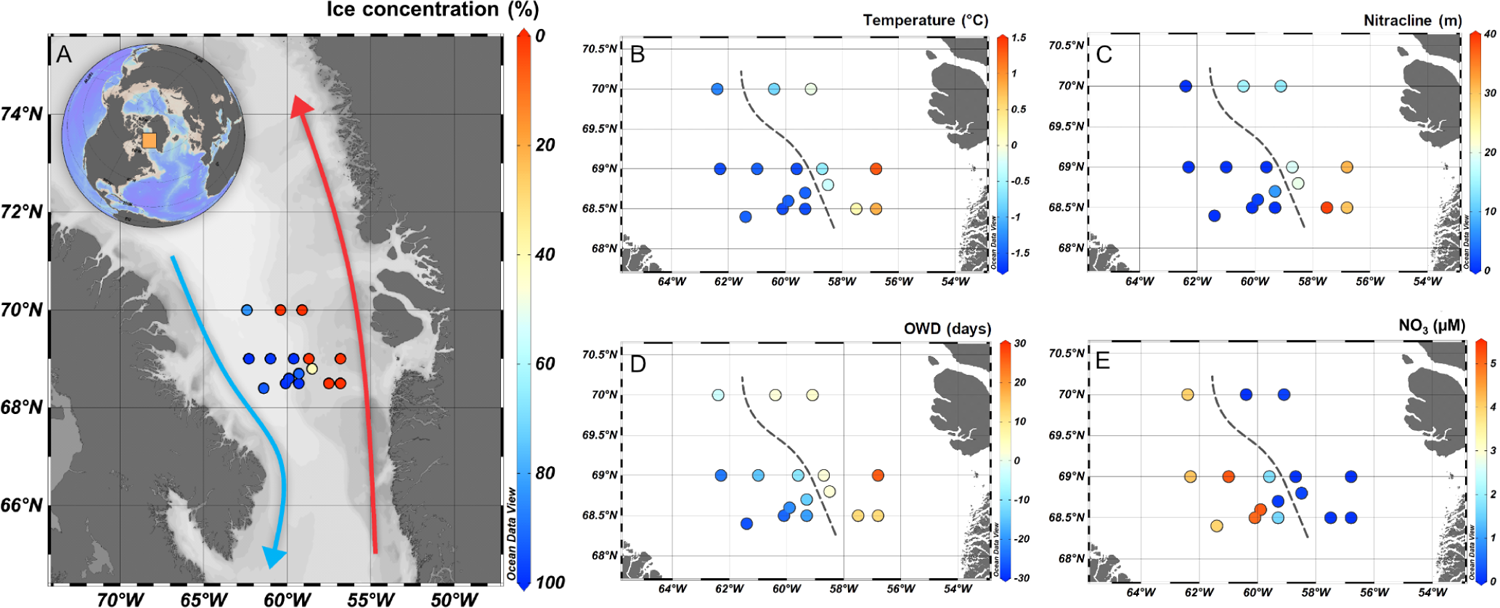
Location of the sampling stations in Baffin Bay and environmental variables. (A) Sampling stations indicating the sea-ice concentration (%); the red arrow represents the warmer West Greenland Current, and the blue arrow represents the Pacific-originated Baffin Current; (B) Temperature (*^◦^*C) in surface; (C) Depth of the nitracline (meters); (D) Open Water Days: amount of days of open water before (positive values) or after (negative values) the sampling day; (E) Nitrate concentration in surface (*µ*M). A dashed line separates sampling stations with more (east) and less (west) than 80% sea-ice cover.

### 18S rRNA V4 PCR amplification and sequencing

The V4 region of the 18S rRNA (about 380 bp) was amplified using the V4 primers TAReuk454FWD1 (forward) and V4 18S Next.Rev (reverse), along with the Illumina Nextera (Illumina, San Diego, CA, USA) 5’ end overhang sequence as described in Piredda et al. (2017). Reaction mixtures in a total of 20 *µ*L were performed using 10 *µ*L of Phusion High-Fidelity PCR Master Mix^®^ 2*×*, 0.3 *µ*M final concentration of each primer, 3% DMSO, 2% BSA and H_2_O. Thermal conditions were as follows: 98*^◦^* C for 5 min, followed by 25 cycles of 98*^◦^* C for 20 s, 52*^◦^* C for 30 s, 72*^◦^* C for 90 s, and a final cycle of 72*^◦^* C for 5 min. Samples were amplified in triplicates and pooled together subsequently in order to minimize the chance of amplification errors. PCR purification was performed using AMPure XP Beads (Beckman Coulter, Brea, CA, USA) following instructions from the manufacturer. DNA quantification and quality check was done using a LabChip GX Touch HT Nucleic Acid Analyzer (PerkinElmer, Waltham, MA, USA). Libraries were prepared as detailed on the Illumina^®^ support website (http://support.illumina.com) with a final concentration of 1 nM and 1% of denaturated PhiX to prevent sequencing errors due to low-diversity libraries. Sequencing was performed using a 2*×*250 bp MiSeq Reagent Kit v2^®^ at the GenoMer platform (Roscoff, France).

### Sequence processing

Sequences were processed using the dada2 (Callahan et al. 2016) package within R (R Core Team 2021). Reads were filtered and trimmed using the filterAndTrim function with the following parameters: truncLen = c(250, 240), trimLeft equal to each primer length (for primer removal), maxN=0, maxEE=c(2, 2), and truncQ=10. Merging of forward and reverse reads with the mergePairs function and chimeric sequences removal with the removeBimeraDenovo function were both performed with default parameters. Resulting ASVs were taxonomically assigned using assignTaxonomy function with PR2 database (Guillou et al. 2013) version 4.14 (https://pr2-database.org/). Samples with less than a total of 3,000 reads were excluded, and the number of reads for each sample was normalized by the median sequencing depth. Autotrophic taxa were selected by filtering-in divisions Chlorophyta, Cryptophyta, Haptophyta, Ochrophyta and Cercozoa. Classes known to comprise only heterotrophic members (Chrysophyceae, Sarcomonadea and Filosa-Thecofilosea) were excluded. Dinoflagellates were not considered because they contain both autotrophic and heterotrophic taxa, and the high number of 18S rRNA gene copies per genome makes them dominate read numbers, obscuring patterns of other autotrophs. Processing script can be found at https://github.com/vaulot/Paper-2021-Vaulot-metapr2/tree/ main/R_processing.

### Environmental data

Environmental variables obtained by other studies during the Green Edge cruise (Lafond et al. 2019; Randelhoff et al. 2019; Saint-Béat et al. 2020) were used here in accordance with our different sectors UI, MIZ and OW. All ancillary physico-chemical and biological data obtained from the Green Edge project used in the present paper is available at http://www.obs-vlfr.fr/proof/php/GREENEDGE/x_datalist_1.php?xxop=greenedge& xxcamp=amundsen as raw data, and at https://www.seanoe.org/data/00487/59892/ (Massicotte et al. 2020) as formatted files, and described in detail by Bruyant et al. (2022) (see Data Availability section). The complete list of variables sampled during the Amundsen Green Edge cruise, the principal investigator responsible for each data set, and the protocols used to obtain and analyze physical, chemical and biological data can be found in Bruyant et al. (2022). Further information on nutrient and pigment analysis can be found in Lafond et al. (2019). Data processing for light transmittance, sea-ice cover and water column stability can be found in Randelhoff et al. (2019).

### Flow cytometry analysis

Autotrophic and heterotrophic cell abundance was measured *in situ* using a BD Accuri*^TM^* C6 flow cytometer as previously described in (Marie et al. 2010). Pico- and nano-phytoplankton abundance was measured on unstained samples with fluorescent beads for parameter normalization (0.95 G Fluoresbrite^®^ Polysciences, Warrington, PA), while heterotrophic cell enumeration was performed using SYBR Green^®^ staining as described in (Marie et al. 1997).

### Data analysis

Sampling stations (Figure 1) were clustered into “Open Water” (OW), “Marginal Ice Zone” (MIZ) and “Under Ice” (UI) on the basis of the temporal dynamics sea ice cover using the parameter “Open Water Days” (OWD). OWD corresponds to how many days a given station had been ice-free before sampling (positive values) or how many days it took for it to become ice-free after sampling (negative values) (see Randelhoff et al. 2019). Stations with OWD > 10 days of open water before sampling day were considered OW, stations with 10 to −10 days were considered within the MIZ, and stations with OWD < −10 days were considered UI (Table 1). The number of sampling points within each sector and size fractions can be found in Table 2.

**Table 1:**
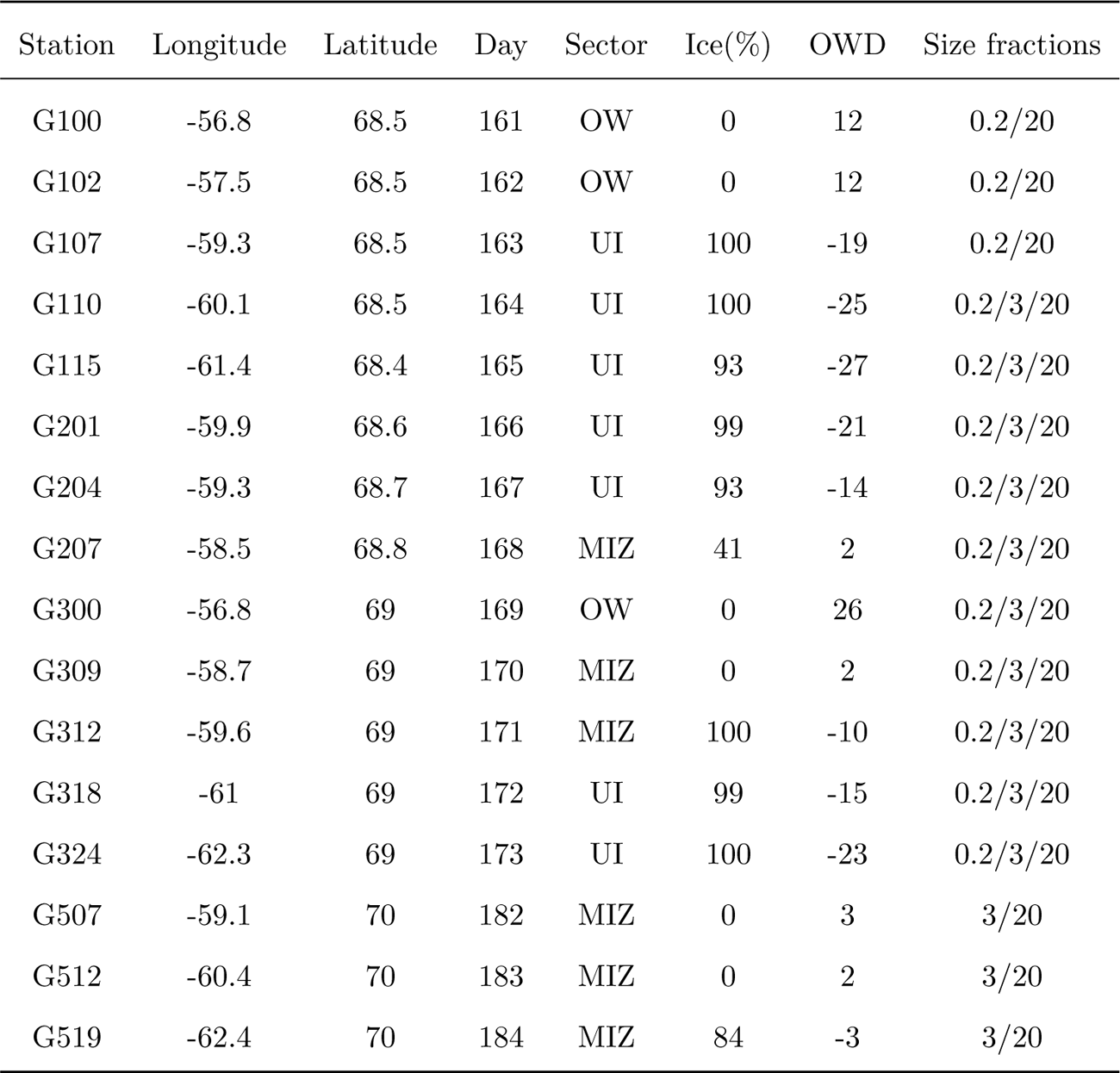
Stations with their geographical coordinates, julian day, the sectors where it belongs (under ice, marginal ice zone or open water), the percentage of ice cover and how many days it has been ice free upon sampling, and size fractions analyzed.

**Table 2:** Number of samples within each sector for each size fraction.

| Sector | 0.2-3 $\mu\text{m}$ | 3-20 $\mu\text{m}$ | > 20 $\mu\text{m}$ |
| --- | --- | --- | --- |
| UI | 40 | 35 | 40 |
| MIZ | 18 | 28 | 34 |
| OW | 16 | 5 | 18 |

Data analysis was performed within R, using the following packages: *phyloseq* (data filtering, heatmaps, alpha diversity) (McMurdie and Holmes 2013), *tidyr* (Wickham et al. 2019), *vegan* (NMDS) (Dixon 2003), *ggplot2* (plotting, Wickham 2016), *ComplexUpset* (upSet graphics, Krassowski 2020). Abundant ASVs for each size fraction were selected by keeping only ASVs which were among the top 90% most abundant sequences in at least one sample. Abundant taxa for the whole community (i.e. considering all size fractions) had to be among the top 90% most abundant sequences in at least 10% of the samples, except for the intersection analysis (upSet graphic), where taxa present in the top 70% of sequences in at least one sample were filtered. This was done with the topf and genefilter_sample functions of *phyloseq*. NMDS analysis was performed using Bray–Curtis distance with the metaMDS function of the package *vegan*, and statistically significant environmental parameters (*p*-value *≤* 0.001) and genera (*p*-value *≤* 0.05) were mapped against it using the function *envfit*. Indicator species analysis (*indicspecies* package, De Cáceres et al. 2010) was performed with abundant taxa (selected as described above) within each size fraction in order to find significant association between taxa and a given sector (or combination of sectors), using the default *IndVal* index as statistic test and 9999 random permutations. Global distribution of ASVs was performed using the metaPR^2^ database (https://shiny.metapr2.org, Vaulot et al. 2022) which contains metabarcodes from 41 public datasets representing more than 4,000 samples distributed over a wide range of ecosystems. ASV sequences from the present study were entered in the “Query” panel, and matching metaPR^2^ ASVs (100% similarity) were displayed in the “Map” panel.

## Results

We sampled the phytoplankton community across the marginal ice zone in Baffin Bay, Arctic, in June and July 2016, to assess changes related to sea-ice melt. The community was sequentially filtered for three size fractions (0.2-3 *µ*m, 3-20 *µ*m and > 20 *µ*m), and sampling stations were classified as Under Ice (UI), Marginal Ice Zone (MIZ) and Open Water (OW) sectors (Tables 1 and 2).

### Physical, chemical and biological variability

Temperature were lower in the Arctic-influenced UI sector and higher in terms of both absolute values and median in the Atlantic-influenced OW sector. Temperature differences between the two sectors were statistically significant (Figure 1 and 2A). Salinity was not significantly different between the two ice-influenced UI and MIZ sectors, with a wider distribution towards less saline sampling points influenced by sea-ice melt. Salinity values were less variable in the OW sector, narrowly ranging between 33.6 and 34 (Figure 2B, Supplementary data S1). Chl fluorescence from the CTD was higher in MIZ and the well-lit OW sector, reaching its peak in the former sector with 14.5 mg.m*^−^*^3^ (Figure 2C). The mixed layer depth (MLD) was significantly different between the UI and OW sectors, being deeper in the UI sector, varying from 27 to 46 m. UI and OW were also distinct from MIZ, with the MLD ranging from 4 to 12 m (Figure 2D). PAR (mol photons*^−^*^2^.d*^−^*^1^) was not significantly different between MIZ and OW, although variability was greater in the MIZ, in keeping with the variable ice cover (Figure 2E).

**Figure 2:**
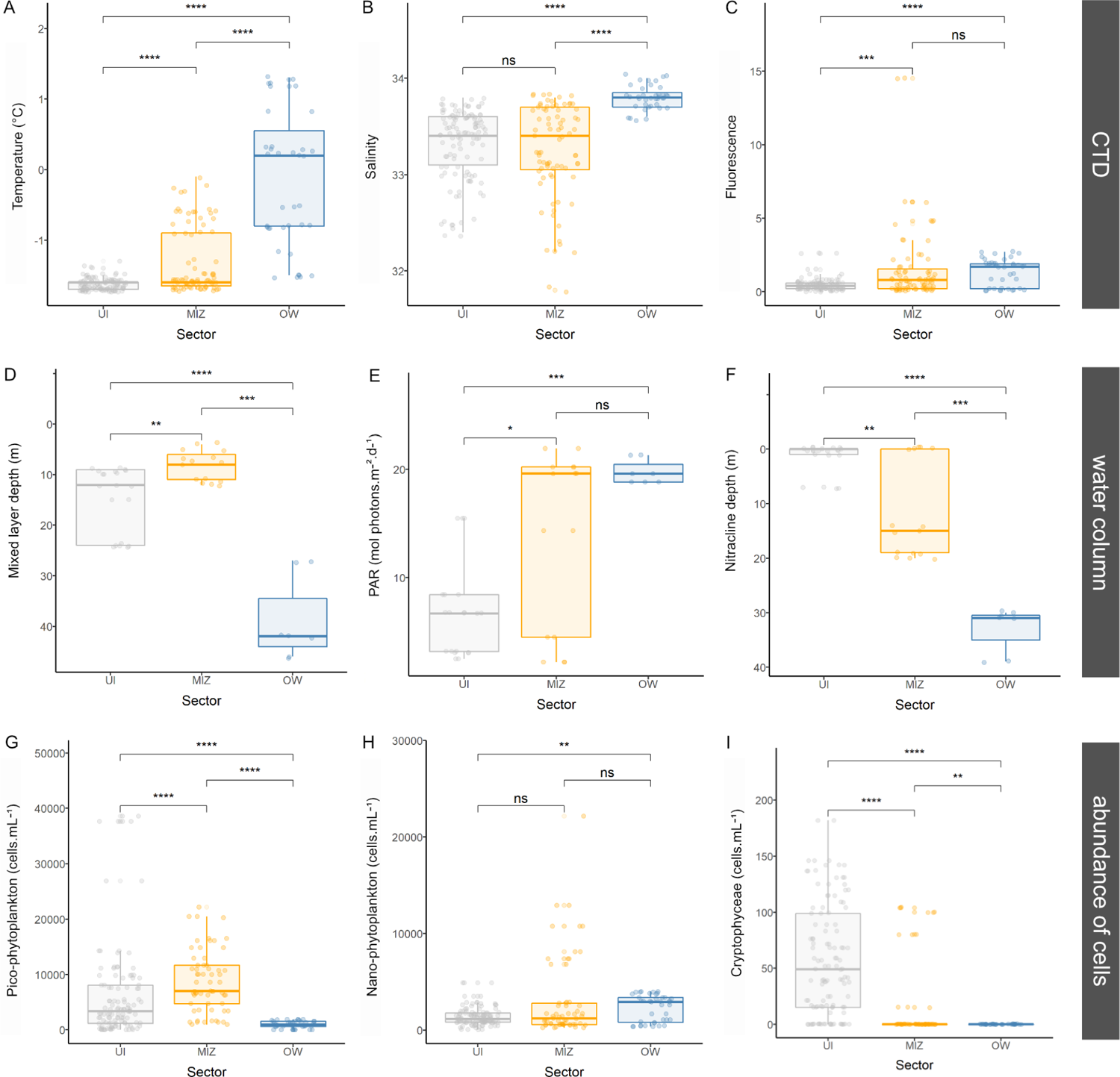
Environmental variables for the three sectors: UI (gray), MIZ (yellow) and OW (blue); (A) temperature (*^◦^*C); (B) salinity; (C) fluorescence; (D) mixed layer depth (m); (E) Photosynthetically Active Radiation at 3 m (mol photons.m^-2^.d*^−^*^1^); (F) nitracline depth (m); (G) pico-phytoplankton abundance (cells.mL*^−^*^1^); (H) nano-phytoplankton abundance (cells.mL*^−^*^1^); (I) Cryptophyceae abundance (cells.mL*^−^*^1^). Number of asterisks represent *p*-value obtained with the Wilcox test as follows: (*) *p ≤* 0.05; (**) *p ≤* 0.01; (***) *p ≤* 0.001; (****) *p ≤* 0.0001; “ns” = not significant.

Nitracline depth was significantly distinct between sectors, in general being deeper in the OW sector and always larger than 30 m. In the UI sector it was never deeper than 8 m, while in the MIZ sector the nitracline depth was variable, with values ranging from 0 and 20 m (Figure 2F). Nutrient concentrations were in general higher in the UI sector, compared to MIZ and UI. MIZ was more similar to OW than UI for all nutrients and ratios measured (Figure S1A-I). Nitrate, phosphate, silica, colored dissolved organic matter (CDOM), urea, particulate organic nitrogen (PON) and carbon (POC) differed significantly between the UI and OW sectors (Figure S1 and S2). Urea concentrations were higher in the UI sector, reaching 1.9 x 10^3^ *µ*M, almost the double the maximum concentration from the other sectors (Figure S1 A). Although ammonium concentrations did not differ significantly between the sectors, values higher than 0.8 *µ*M were only found in the UI sector, up to 7.7 *µ*M (Figure S1 B). Ammonium assimilation and regeneration were significantly different between the different sectors, with higher median values found in the MIZ sector (Figure S2 G-H), while urea assimilation decreased in the UI sector and nitrate assimilation was somewhat even among all sectors (Figure S2 I-J). DON and primary production were higher in the MIZ sector, although the highest values from the latter were obtained in the UI sector, up to 88 *µ*gC.L*^−^*^1^.day*^−^*^1^ (Figure S2 L).

### Phytoplankton abundance

Phytoplankton abundance measured by flow cytometry revealed different distribution patterns between pico (0.2-3 *µ*m) and nano (3-20 *µ*m) size fractions (Figure 2G-H). Pico-phytoplankton abundance was greatest in the UI sector (up to 39 x 10^3^ cells.mL*^−^*^1^), and lowest values in the OW sector (0.95 x 10^3^ cells.mL*^−^*^1^ on average) (Figure 2G). Differences between sectors were highly significant for the smallest size fraction, in contrast to nanophytoplankton, where only the extremes UI and OW differed significantly. Nano-phytoplankton abundance was the highest in the MIZ sector (up to 22 x 10^3^ cells.mL*^−^*^1^), although the median was the highest in OW (Figure 2H). FCM cryptophyte abundance differed significantly among all sectors, with much higher values in UI (up to 182 cells.mL*^−^*^1^) than in MIZ and especially in OW, where they were virtually absent (Figure 2I). Pico- and nano-phytoplankton abundance was in general higher in surface and decreased with depth in UI, while in OW the pattern was the inverse (Figure 3). Within the MIZ pico-phytoplankton abundance was higher in surface and subsurface, while nano-phytoplankton peaked in deeper samples. Cryptophyceae abundance in UI sector was in general higher in surface/subsurface, but with some abundance peaks in deeper samples (Figure 3).

**Figure 3:**
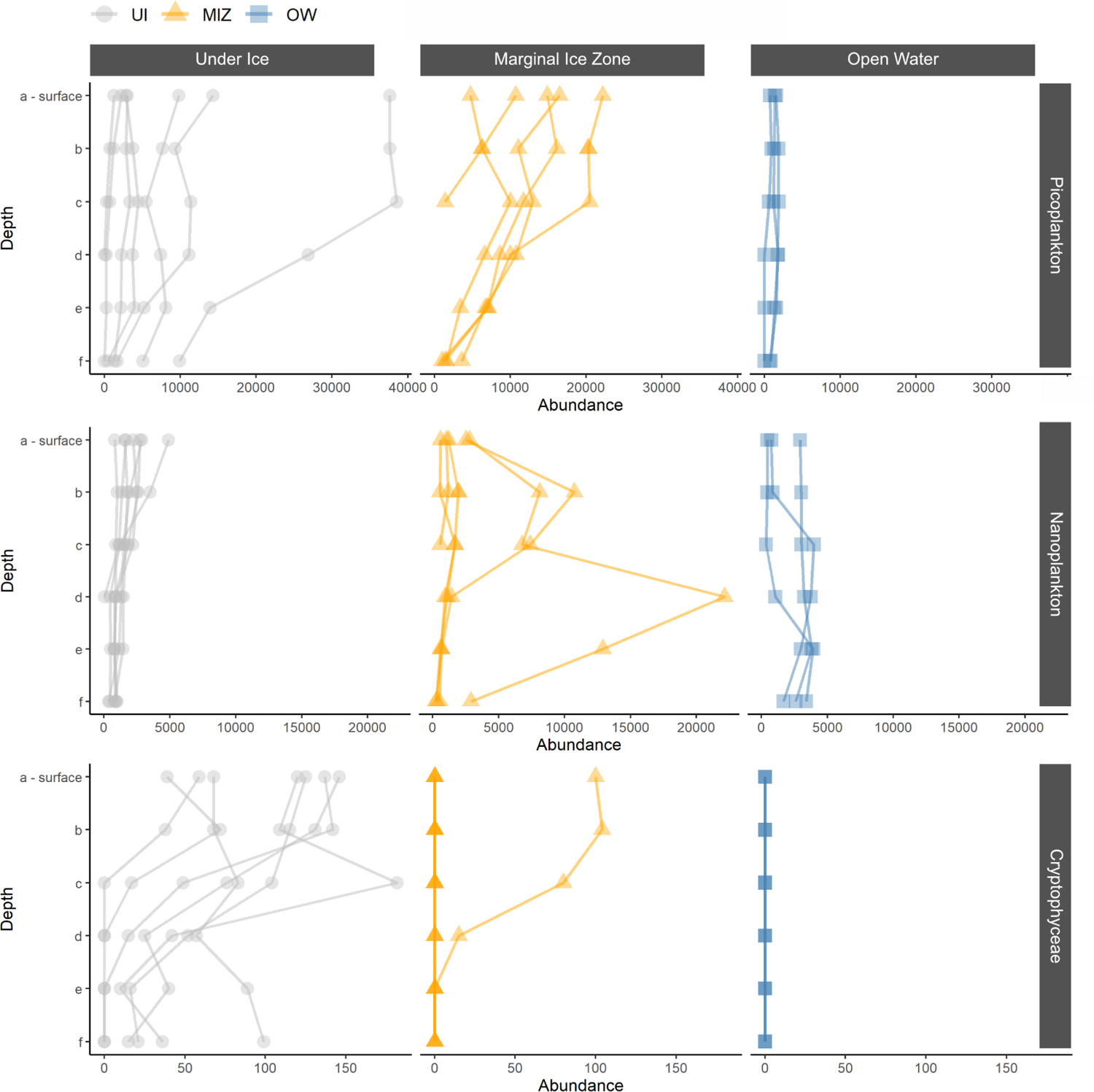
Abundance (cells.mL*^−^*^1^) measured by FCM of pico-phytoplankton (top panels), nano-phytoplankton (middle panels) and Cryptophyceae (lower panels) according to depth, divided between the three sectors: UI (gray), MIZ (yellow) and OW (blue).

### Phytoplankton diversity at the division and genus level

Diversity patterns at the division level had a marked difference between sectors, especially for the smaller (0.2-3 and 3-20 *µ*m) size fractions. The 0.2-3 *µ*m size fraction was mostly dominated by Chlorophyta in the ice-associated (UI+MIZ) sectors throughout the water column, with an important share of Cryptophyta and Ochrophyta, while in the OW sector Haptophyta reads were predominant, especially in deeper samples (Figure S3). Differences in diversity between sectors at the division level in the 3-20 *µ*m size fraction were less marked, although there was in general an increase in Haptophyta towards the MIZ and OW sectors. This size fraction separation was also marked with more Ochrophyta that dominated the OW sector in surface samples (Figure S3). In the > 20 *µ*m size fraction, the Ochrophyta abundance dominated the three sectors with only a small increase in Haptophyta relative abundance towards MIZ and OW sectors (Figure S3).

**0.2-3** *µ***m**. At the genus level, diversity in the 0.2-3 *µ*m size fraction did not differ greatly across sectors from that observed at the division level, since the two most abundant divisions, Chlorophyta and Haptophyta, were dominated by the genera *Micromonas* and *Phaeocystis*, respectively (Figure 4). Within Mamiellophyceae, *Bathycoccus* and *Mantoniella* were mainly detected in ice-associated sectors, the former with higher relative abundances in the deeper samples and the latter in surface samples. Cryptophyta was mainly dominated by *Falcomonas* in UI and MIZ, and by *Teleaulax* in the OW sector. Although not very abundant, Bacillariophyceae were extremely diverse in ice-associated sectors (Figure 4). It is important to note that the decrease in *Micromonas* towards the OW sector is corroborated by the large drop in pico-phytoplankton cells abundance within this sector (Figure 2G).

**Figure 4:**
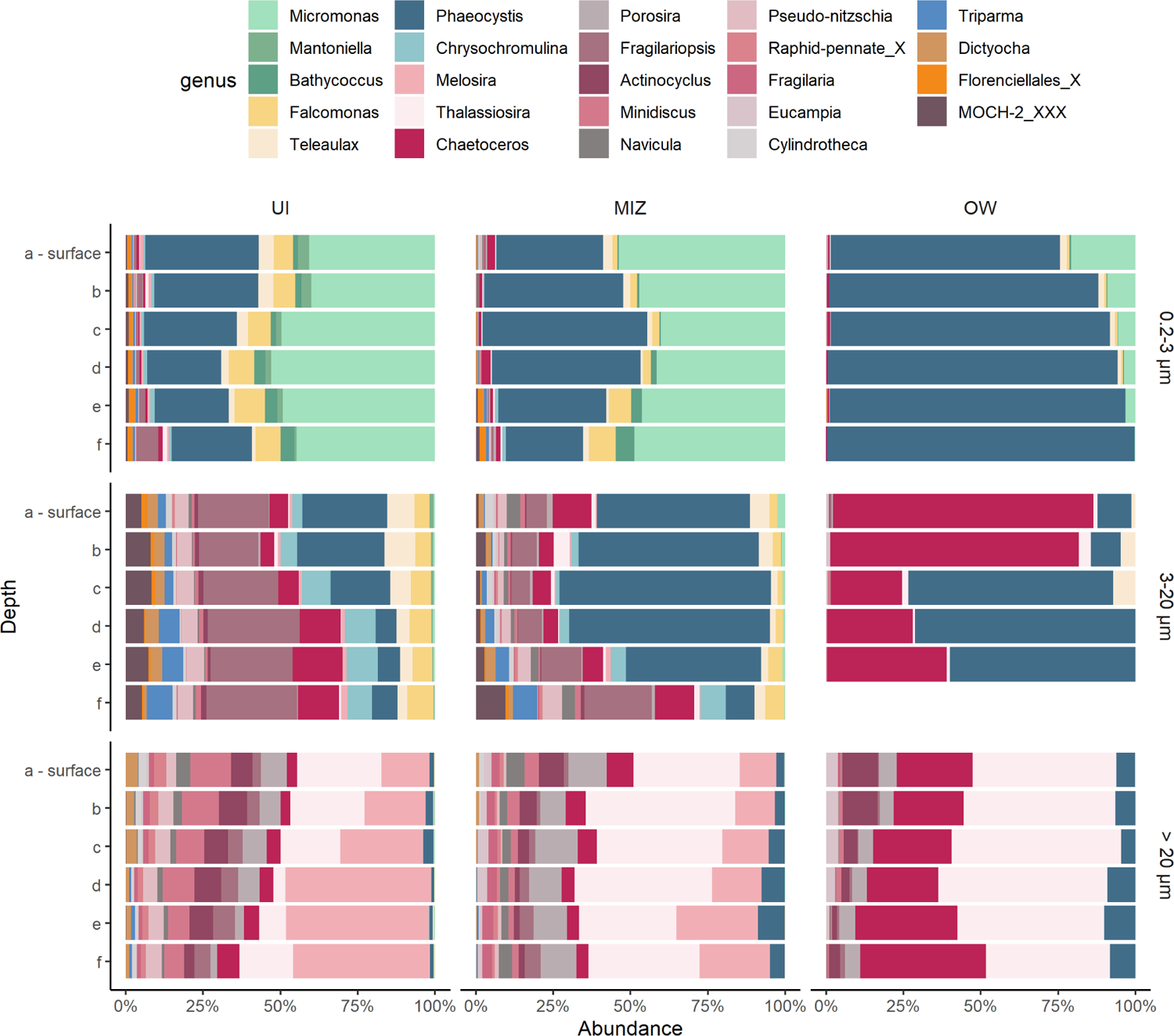
Relative abundance of reads at the genus level between sectors and size fractions. UI: Under Ice; MIZ: Marginal Ice Zone; OW: Open Water; letters on the y-axis refer to the depth level where “a” corresponds to the surface and “f” to the deepest sample depth, usually between 40 m and 60 m depth.

**3-20** *µ***m**. Mamiellophyceae were nearly absent in the 3-20 *µ*m size fraction, except for a small contribution to surface samples in UI and MIZ. A higher contribution of *Chrysochromulina* within Haptophyta was observed in ice-associated sectors, especially in deeper samples in the UI sector (Figure 4). A higher abundance of *Teleaulax* relative to *Falcomonas* was observed when comparing the 3-20 *µ*m to the 0.2-3 *µ*m size fraction, always more present in surface than deeper samples, including in the OW sector. As observed in the smallest size fraction, Ochrophyta were fairly diverse in ice-associated sectors, with representatives of Bacillariophyceae, Bolidophyceae, Dictyochophyceae, and Marine Ochrophyta (MOCH-2). *Chaetoceros* was dominant in the OW sector, especially in surface samples, with a small contribution of *Thalassiosira*.

**> 20** *µ***m**. There was a decrease in non-diatom Ochrophyta representatives in the > 20 *µ*m size fraction, although *Dictyocha* and *Triparma* were still present in ice-associated sectors, the former mostly in surface and the latter in deeper samples (Figure 4). With respect to diatoms, there was an increase in *Porosira*, *Actinocyclus*, and especially *Thalassiosira* in all the sectors in comparison with other size fractions, and in *Melosira* relative abundance in ice-associated sectors.

NMDS analysis revealed that samples clustered according to size fractions along the first axis and sectors along the second axis. UI and MIZ were associated with higher nutrient concentration and Cryptophyceae cell abundance, whereas OW sector presented higher temperatures and use of alternative source of nitrogen, such as urea and ammonium, indicating the importance of regenerated production (Figure 5A). Statistically significant genera had a distribution linked to both sectors and size fractions. For example, larger size fractions from ice-associated samples were correlated with pennate diatoms such as *Pseudo-nitzschia* and *Cylindrotheca*, whereas smaller size fractions for the same samples were correlated with *Falcomonas*, *Bathycoccus* and *Micromonas*. Larger size fractions from the OW sector were associated with centric diatoms such as *Thalassiosira*, *Chaetoceros* and *Eucampia*, and smaller size fractions with *Phaeocystis*.

**Figure 5:**
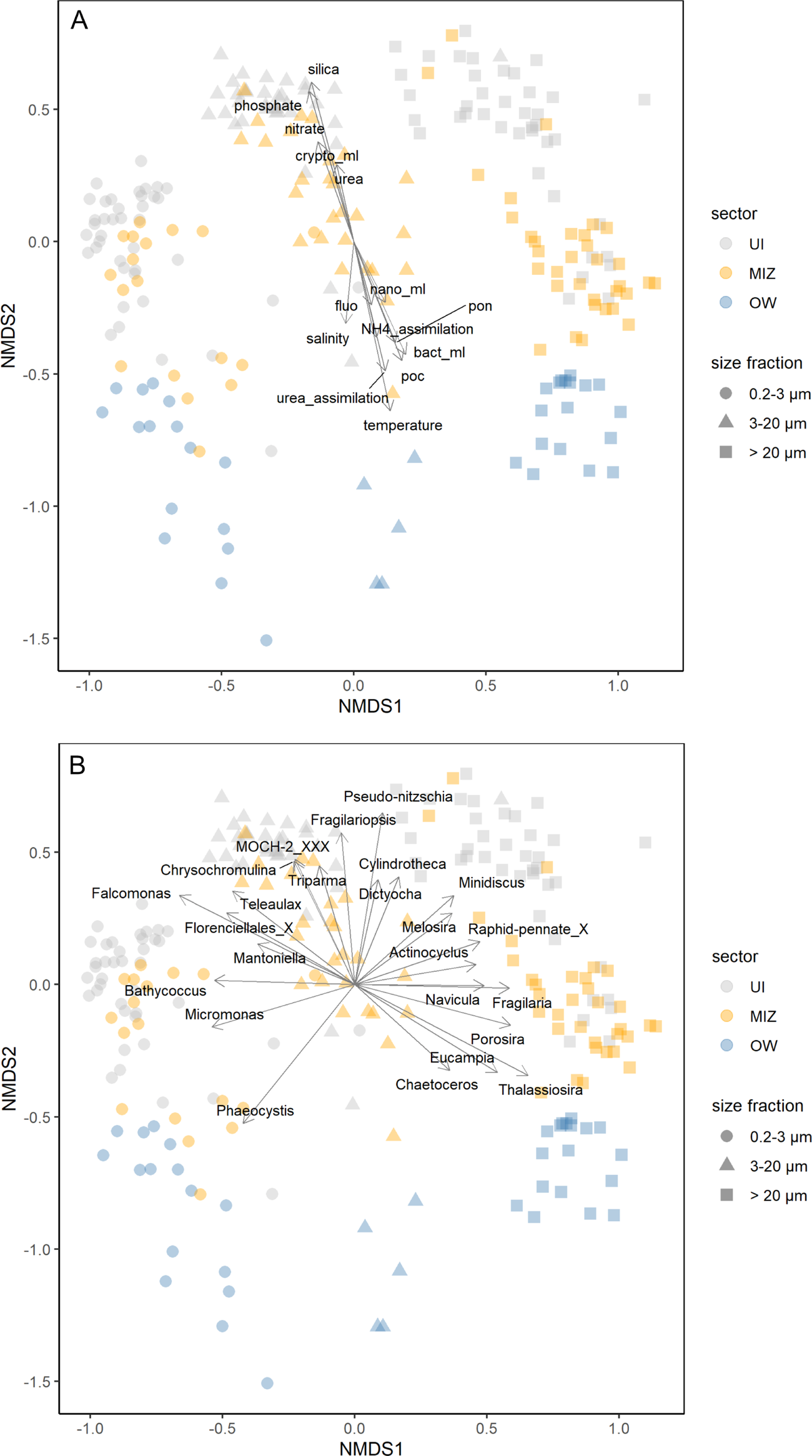
Non-metric multidimensional scaling (NMDS) analysis using Bray-Curtis dissimilarities of the phytoplankton community composition; only statistically significant (A) environmental parameters (*p*-value = 0.001) and (B) genera (*p*-value = 0.05) were plotted against ordination. Parameters: ammonium assimilation (NH4_assimilation), bacteria abundance (bact_ml), Cryptophyceae abundance (crypto_ml), fluorescence (fluo), nano-phytoplankton abundance (nano_ml), nitrates, silica, particulate organic carbon (poc), particulate organic nitrogen (pon), phosphate, salinity, temperature, urea, and urea assimilation (urea_assimilation). Stress: 0.11.

### Phytoplankton microdiversity

Taxa grouped by genera mask variability at the species and ASV level. Looking at all the genera with more than two ASVs in the whole dataset, the ASV-level distribution of taxa yields potential information on niche-preference. In general, most genera had more ice-associated ASVs compared to OW (Table 3). Interestingly, several low-abundance taxa harbored high numbers of ice-associated ASVs, for example, the Dictyochophyceae genus *Pseudochattonella* and the environmental clade 2 of Bolidophyceae. Two groups presented a surprisingly large number of ASVs: the B clade of Dolichomastigaceae (Mamiellophyceae) with a total of 29 ASVs, and the centric diatom genus *Chaetoceros* with 35 ASVs (Table 3).

**Table 3:**
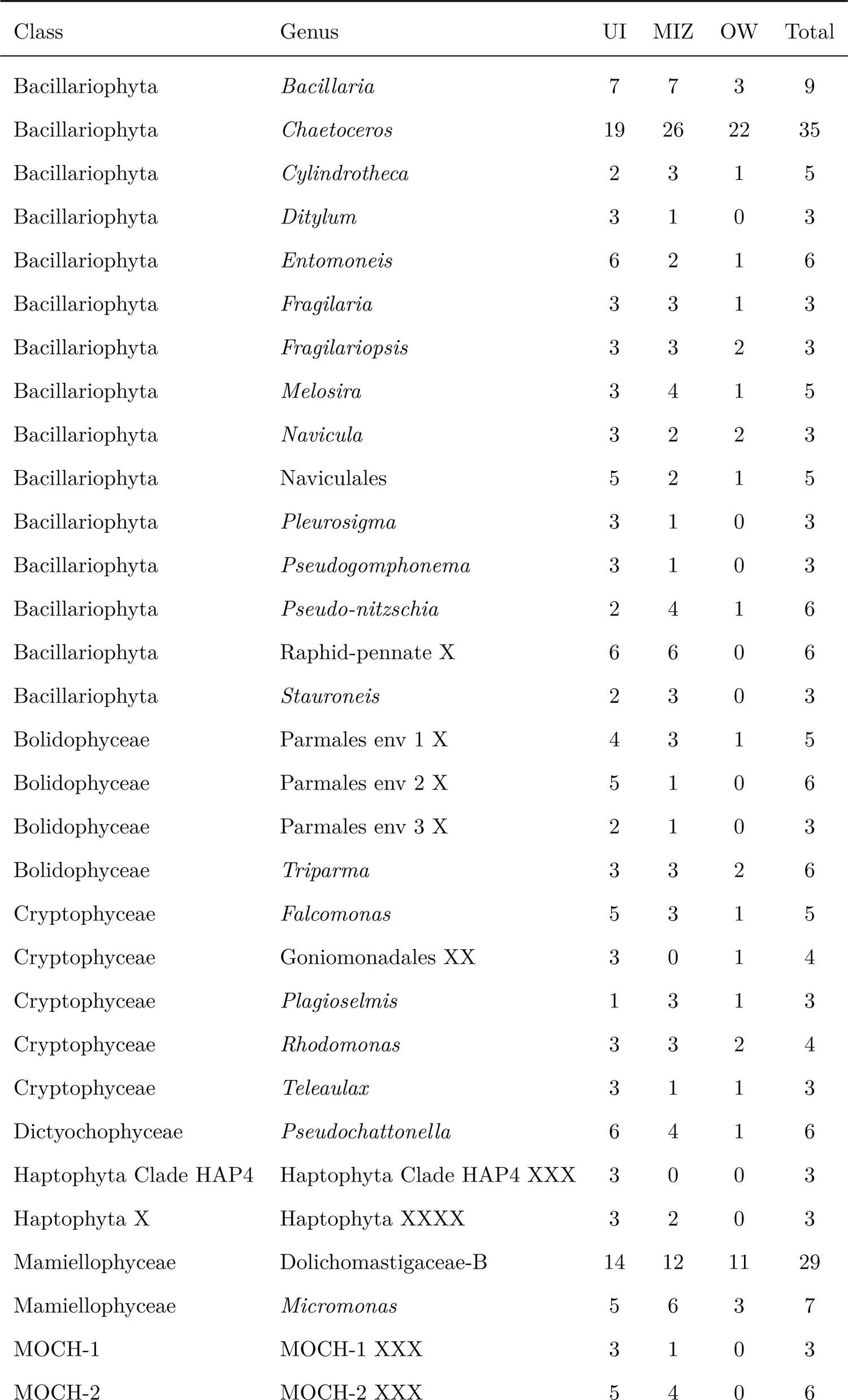
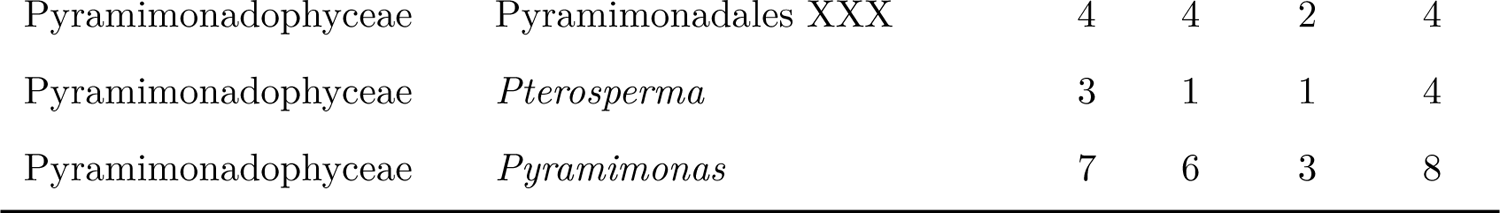
Number of ASVs by phytoplankton genera present in each sector; only genera with more than 2 ASVs in the whole dataset were taken into account. Note that taxa not assigned to the genus level might contain more than one genus.

Alpha diversity indices indicate that, in general, diversity was higher in the smallest size fraction in the UI and MIZ sectors, and decreased towards bigger size fractions. However, the Simpson index was lowest in the 3-20 *µ*m size fraction and the highest in the > 20 *µ*m size fraction (Figure S4). When taking into consideration only the most abundant ASVs (ranking among the top 70% most abundant sequences in at least one sample), 41 ASVs were shared between all sectors, 16 were exclusive to the UI and MIZ sectors, and 2 were exclusive from the UI sector, while none was exclusive to the OW sector or to the MIZ and OW taken together (Figure S5).

From the ten ASVs exclusive from ice-associated sectors, four were diatoms, three pennate and one centric diatom, which was also the most abundant, *M. arctica* (ASV_0025). Three of these ASVs were Haptophyta, two assigned as *Chrysochromulina* and one to *Phaeocystis cordata*, a species described from the Mediterranean Sea (Zingone et al. 1999). The other three ASVs were the only representatives of their classes: the uncultivated MOCH-2 (ASV_0061), the Cryptophyceae *F. daucoides* (ASV_0055), and the Mamiellophyceae *Mantoniella squamata* (ASV_0104) (Figure S5).

### Indicator ASVs

In order to find patterns of taxa distribution that could be used as ecological indicators of niche preferences, we analyzed ASVs distribution on each sector and group of sectors using statistical indices described by De Cáceres et al., (2010).

**0.2-3** *µ***m**. The indicator species analysis identified 39 ASVs that were representatives of one sector or a group of sectors within the 0.2-3 *µ*m size fraction, 30 of them related to UI (10), MIZ (2), or the UI+MIZ sector group (18) (Table 4). Among the highly significant taxa within the UI sector (*p*-value < 0.001) there were four Ochrophyta, three diatoms (*M. arctica*, *Fragilariopsis cylindrus* and *Bacillaria paxillifer*), and one Pelagophyceae (assigned to the genus *Ankylochrysis*). Thirteen AVSs were highly correlated to the MIZ+UI sector grouping, including two Mamiellophyceae (*B. prasinos* and *Micromonas commoda* A2), two cryptophytes, both assigned to *F. daucoides*, seven non-diatom Ochrophyta (from the classes MOCH-2, Bolidophyceae, Dictyochophyceae, and Pelagophyceae), and two diatoms (*Pseudo-nitzschia seriata* and *Chaetoceros neogracilis*). A *M. polaris* ASV (ASV_0154) was also considered indicator of the MIZ+UI site group, with a *p*-value of 0.002. Taxa indicators of the OW sector (4) included three undescribed Dictyochophyceae (all assigned to *Pedinellales* sp.), and one undescribed Dolichomastigaceae from clade B, while the three taxa indicators of the MIZ+OW sector group comprised two centric diatoms (*Porosira glacialis* and *Chaetoceros decipiens*) and one undescribed Prymnesiophyceae.

**Table 4:** Indicator ASVs with their taxonomic assignation for each sector or group of sectors, divided by size fraction. “A” represents the positive predictive power of the ASV, or the probability of a sampling site being a member of the sector or group of sectors when the ASV appears in that site. “B” represents how often one ASV is found in sampling sites of the sector or group of sectors. The value of the correlation (stat) and the statistical significance of the association (*p*-value) are also shown.

| Size fraction | Sectors | ASVs | Class | Species | A | B | Stat | p.value | Sign |
| --- | --- | --- | --- | --- | --- | --- | --- | --- | --- |
| 0.2-3 $\mu\text{m}$ | UI | asv_023_00009 | Bacillariophyta | Melosira arctica | 0.98 | 0.7 | 0.83 | 1e-04 | *** |
| 0.2-3 $\mu\text{m}$ | UI | asv_023_00015 | Bacillariophyta | Fragilariopsis cylindrus | 0.93 | 0.75 | 0.83 | 1e-04 | *** |
| 0.2-3 $\mu\text{m}$ | UI | asv_023_00104 | Mamiellophyceae | Mantoniella squamata | 1 | 0.48 | 0.69 | 2e-04 | *** |
| 0.2-3 $\mu\text{m}$ | UI | asv_023_00125 | Prymnesiophyceae | Phaeocystis sp. | 0.93 | 0.75 | 0.84 | 1e-04 | *** |
| 0.2-3 $\mu\text{m}$ | UI | asv_023_00171 | Bacillariophyta | Bacillaria paxillifer | 0.97 | 0.68 | 0.81 | 1e-04 | *** |
| 0.2-3 $\mu\text{m}$ | UI | asv_023_00244 | Pyramimonadales | Pterosperma sp. | 1 | 0.42 | 0.65 | 5e-04 | *** |
| 0.2-3 $\mu\text{m}$ | UI | asv_023_00311 | Pelagophyceae | Ankylochrysis sp. | 0.96 | 0.48 | 0.67 | 4e-04 | *** |
| 0.2-3 $\mu\text{m}$ | UI | asv_023_00384 | Mamiellophyceae | Dolichomastigaceae-B sp. | 1 | 0.32 | 0.57 | 0.0031 | ** |
| 0.2-3 $\mu\text{m}$ | UI | asv_023_00239 | Prymnesiophyceae | Chrysochromulina sp. | 1 | 0.22 | 0.47 | 0.0189 | * |
| 0.2-3 $\mu\text{m}$ | UI | asv_023_00381 | Haptophyta_Clade_HAP4 | Haptophyta Clade HAP4 XXX sp. | 1 | 0.28 | 0.52 | 0.0107 | * |
| 0.2-3 $\mu\text{m}$ | MIZ | asv_023_00008 | Bacillariophyta | Thalassiosira antarctica | 0.52 | 0.78 | 0.63 | 0.0186 | * |
| 0.2-3 $\mu\text{m}$ | MIZ | asv_023_00049 | Bacillariophyta | Navicula sp. | 0.74 | 0.39 | 0.54 | 0.0067 | ** |
| 0.2-3 $\mu\text{m}$ | OW | asv_023_00421 | Mamiellophyceae | Dolichomastigaceae-B sp. | 0.89 | 0.44 | 0.62 | 2e-04 | *** |
| 0.2-3 $\mu\text{m}$ | OW | asv_023_00469 | Dictyochophyceae | Pedinellales X sp. | 0.98 | 0.81 | 0.89 | 1e-04 | *** |
| 0.2-3 $\mu\text{m}$ | OW | asv_023_00593 | Dictyochophyceae | Pedinellales X sp. | 1 | 0.5 | 0.71 | 1e-04 | *** |
| 0.2-3 $\mu\text{m}$ | OW | asv_023_00731 | Dictyochophyceae | Pedinellales X sp. | 1 | 0.31 | 0.56 | 3e-04 | *** |
| 0.2-3 $\mu\text{m}$ | OW | asv_023_00223 | Prymnesiophyceae | Prymnesiophyceae Clade F XX sp. | 0.89 | 0.19 | 0.41 | 0.0149 | * |
| 0.2-3 $\mu\text{m}$ | OW | asv_023_00949 | Mamiellophyceae | Dolichomastigaceae-B sp. | 1 | 0.12 | 0.35 | 0.0447 | * |
| 0.2-3 $\mu\text{m}$ | MIZ+UI | asv_023_00086 | Prymnesiophyceae | Chrysochromulina sp. | 1 | 0.33 | 0.57 | 0.0262 | * |
| 0.2-3 $\mu\text{m}$ | MIZ+UI | asv_023_00041 | Cryptophyceae | Falcomonas daucoides | 0.95 | 0.91 | 0.93 | 1e-04 | *** |
| 0.2-3 $\mu\text{m}$ | MIZ+UI | asv_023_00046 | Bacillariophyta | Pseudo-nitzschia seriata | 1 | 0.59 | 0.77 | 2e-04 | *** |
| 0.2-3 $\mu\text{m}$ | MIZ+UI | asv_023_00048 | Bacillariophyta | Chaetoceros neogracilis | 0.97 | 0.79 | 0.88 | 1e-04 | *** |
| 0.2-3 $\mu\text{m}$ | MIZ+UI | asv_023_00055 | Cryptophyceae | Falcomonas daucoides | 1 | 0.57 | 0.75 | 8e-04 | *** |
| 0.2-3 $\mu\text{m}$ | MIZ+UI | asv_023_00061 | MOCH-2 | MOCH-2 XXX sp. | 1 | 0.78 | 0.88 | 1e-04 | *** |
| 0.2-3 $\mu\text{m}$ | MIZ+UI | asv_023_00073 | Bolidophyceae | Triparma laevis clade | 0.97 | 0.76 | 0.86 | 2e-04 | *** |
| 0.2-3 $\mu\text{m}$ | MIZ+UI | asv_023_00075 | Dictyochophyceae | Dictyocha sp.eculum | 1 | 0.62 | 0.79 | 2e-04 | *** |
| 0.2-3 $\mu\text{m}$ | MIZ+UI | asv_023_00081 | Mamiellophyceae | Bathycoccus prasinos | 0.96 | 0.91 | 0.94 | 1e-04 | *** |
| 0.2-3 $\mu\text{m}$ | MIZ+UI | asv_023_00192 | Prymnesiophyceae | Haptolina sp. | 0.94 | 0.45 | 0.65 | 0.0444 | * |
| 0.2-3 $\mu\text{m}$ | MIZ+UI | asv_023_00105 | Prymnesiophyceae | Phaeocystis cordata | 1 | 0.5 | 0.71 | 0.0013 | ** |
| 0.2-3 $\mu\text{m}$ | MIZ+UI | asv_023_00114 | Dictyochophyceae | Florenciellales X sp. | 0.98 | 0.66 | 0.8 | 2e-04 | *** |
| 0.2-3 $\mu\text{m}$ | MIZ+UI | asv_023_00133 | Pelagophyceae | Pelagomonas calceolata | 0.98 | 0.83 | 0.9 | 1e-04 | *** |
| 0.2-3 $\mu\text{m}$ | MIZ+UI | asv_023_00151 | Dictyochophyceae | Florenciella parvula | 0.97 | 0.83 | 0.9 | 1e-04 | *** |
| 0.2-3 $\mu\text{m}$ | MIZ+UI | asv_023_00267 | Mamiellophyceae | Dolichomastigaceae-B sp. | 1 | 0.29 | 0.54 | 0.0393 | * |
| 0.2-3 $\mu\text{m}$ | MIZ+UI | asv_023_00154 | Mamiellophyceae | Micromonas polaris | 1 | 0.48 | 0.7 | 0.0017 | ** |
| 0.2-3 $\mu\text{m}$ | MIZ+UI | asv_023_00235 | Mamiellophyceae | Micromonas commoda A2 | 0.96 | 0.9 | 0.93 | 1e-04 | *** |
| 0.2-3 $\mu\text{m}$ | MIZ+UI | asv_023_00236 | Dictyochophyceae | Pseudochattonella sp. | 1 | 0.76 | 0.87 | 1e-04 | *** |
| 0.2-3 $\mu\text{m}$ | MIZ+OW | asv_023_00016 | Bacillariophyta | Porosira glacialis | 0.87 | 0.53 | 0.68 | 9e-04 | *** |
| 0.2-3 $\mu\text{m}$ | MIZ+OW | asv_023_00136 | Bacillariophyta | Chaetoceros decipiens | 0.91 | 0.88 | 0.9 | 1e-04 | *** |
| 0.2-3 $\mu\text{m}$ | MIZ+OW | asv_023_00247 | Prymnesiophyceae | Prymnesiophyceae Clade E XX sp. | 0.93 | 0.5 | 0.68 | 5e-04 | *** |
| 3-20 $\mu\text{m}$ | UI | asv_023_00171 | Bacillariophyta | Bacillaria paxillifer | 0.94 | 0.6 | 0.75 | 0.0031 | ** |
| 3-20 $\mu\text{m}$ | UI | asv_023_00244 | Pyramimonadales | Pterosperma sp. | 1 | 0.49 | 0.7 | 0.0095 | ** |
| 3-20 $\mu\text{m}$ | UI | asv_023_00104 | Mamiellophyceae | Mantoniella squamata | 0.94 | 0.46 | 0.66 | 0.0175 | * |
| 3-20 $\mu\text{m}$ | UI | asv_023_00281 | Cryptophyceae | Cryptomonadales XX sp. | 0.87 | 0.57 | 0.71 | 0.0251 | * |
| 3-20 $\mu\text{m}$ | UI | asv_023_00239 | Prymnesiophyceae | Chrysochromulina sp. | 0.94 | 0.4 | 0.61 | 0.0327 | * |
| 3-20 $\mu\text{m}$ | UI | asv_023_00311 | Pelagophyceae | Ankylochrysis sp. | 0.86 | 0.46 | 0.63 | 0.0421 | * |
| 3-20 $\mu\text{m}$ | MIZ | asv_023_00064 | Bacillariophyta | Fragilaria sp. | 0.89 | 0.44 | 0.63 | 0.0267 | * |
| 3-20 $\mu\text{m}$ | MIZ | asv_023_00347 | MOCH-2 | MOCH-2 XXX sp. | 0.89 | 0.59 | 0.73 | 0.0127 | * |

Table 4: (continued)
| Size fraction | Sectors | ASVs | Class | Species | A | B | Stat | p.value | Sign |
| --- | --- | --- | --- | --- | --- | --- | --- | --- | --- |
| 3-20 $\mu\text{m}$ | MIZ | asv_023_00016 | Bacillariophyta | Porosira glacialis | 0.82 | 0.74 | 0.78 | 0.0047 | ** |
| 3-20 $\mu\text{m}$ | MIZ | asv_023_00277 | Bacillariophyta | Naviculales sp. | 0.93 | 0.37 | 0.59 | 0.0488 | * |
| 3-20 $\mu\text{m}$ | MIZ | asv_023_00663 | Filosa-Imbricatea | Novel-clade-2 X sp. | 1 | 0.19 | 0.43 | 0.0438 | * |
| 3-20 $\mu\text{m}$ | MIZ | asv_023_00008 | Bacillariophyta | Thalassiosira antarctica | 0.77 | 0.74 | 0.76 | 0.0166 | * |
| 3-20 $\mu\text{m}$ | MIZ | asv_023_00049 | Bacillariophyta | Navicula sp. | 0.82 | 0.89 | 0.85 | 6e-04 | *** |
| 3-20 $\mu\text{m}$ | OW | asv_023_00320 | Bacillariophyta | Navicula sp. | 0.54 | 0.8 | 0.66 | 0.0141 | * |
| 3-20 $\mu\text{m}$ | OW | asv_023_00057 | Bacillariophyta | Thalassiosira sp. | 0.92 | 0.8 | 0.86 | 1e-04 | *** |
| 3-20 $\mu\text{m}$ | OW | asv_023_00177 | Bacillariophyta | Chaetoceros rostratus | 0.83 | 1 | 0.91 | 1e-04 | *** |
| 3-20 $\mu\text{m}$ | MIZ+UI | asv_023_00009 | Bacillariophyta | Melosira arctica | 1 | 0.69 | 0.83 | 0.0045 | ** |
| 3-20 $\mu\text{m}$ | MIZ+UI | asv_023_00015 | Bacillariophyta | Fragilariopsis cylindrus | 1 | 0.98 | 0.99 | 1e-04 | *** |
| 3-20 $\mu\text{m}$ | MIZ+UI | asv_023_00041 | Cryptophyceae | Falcomonas daucoides | 1 | 0.81 | 0.9 | 8e-04 | *** |
| 3-20 $\mu\text{m}$ | MIZ+UI | asv_023_00046 | Bacillariophyta | Pseudo-nitzschia seriata | 1 | 0.87 | 0.93 | 2e-04 | *** |
| 3-20 $\mu\text{m}$ | MIZ+UI | asv_023_00048 | Bacillariophyta | Chaetoceros neogracilis | 0.99 | 0.98 | 0.99 | 1e-04 | *** |
| 3-20 $\mu\text{m}$ | MIZ+UI | asv_023_00022 | Bacillariophyta | Actinocyclus curvatulus | 0.98 | 0.77 | 0.87 | 0.0126 | * |
| 3-20 $\mu\text{m}$ | MIZ+UI | asv_023_00099 | Bacillariophyta | Cylindrotheca closterium | 1 | 0.65 | 0.8 | 0.0143 | * |
| 3-20 $\mu\text{m}$ | MIZ+UI | asv_023_00061 | MOCH-2 | MOCH-2 XXX sp. | 1 | 0.94 | 0.97 | 2e-04 | *** |
| 3-20 $\mu\text{m}$ | MIZ+UI | asv_023_00073 | Bolidophyceae | Triparma laevis clade | 1 | 0.89 | 0.94 | 1e-04 | *** |
| 3-20 $\mu\text{m}$ | MIZ+UI | asv_023_00075 | Dictyochophyceae | Dictyocha speculum | 1 | 0.79 | 0.89 | 9e-04 | *** |
| 3-20 $\mu\text{m}$ | MIZ+UI | asv_023_00109 | Bacillariophyta | Fragilariopsis sublineata | 1 | 0.61 | 0.78 | 0.0192 | * |
| 3-20 $\mu\text{m}$ | MIZ+UI | asv_023_00003 | Mamiellophyceae | Micromonas polaris | 1 | 0.61 | 0.78 | 0.0189 | * |
| 3-20 $\mu\text{m}$ | MIZ+UI | asv_023_00114 | Dictyochophyceae | Florenciellales X sp. | 1 | 0.73 | 0.85 | 0.0046 | ** |
| 3-20 $\mu\text{m}$ | MIZ+UI | asv_023_00105 | Prymnesiophyceae | Phaeocystis cordata | 1 | 0.58 | 0.76 | 0.0289 | * |
| 3-20 $\mu\text{m}$ | MIZ+UI | asv_023_00125 | Prymnesiophyceae | Phaeocystis sp. | 1 | 0.65 | 0.8 | 0.0215 | * |
| 3-20 $\mu\text{m}$ | MIZ+UI | asv_023_00149 | Cryptophyceae | Plagioselmis prolonga | 0.99 | 0.56 | 0.75 | 0.0364 | * |
| 3-20 $\mu\text{m}$ | MIZ+OW | asv_023_00079 | Bacillariophyta | Eucampia sp. | 0.93 | 0.75 | 0.83 | 5e-04 | *** |
| 3-20 $\mu\text{m}$ | MIZ+OW | asv_023_00136 | Bacillariophyta | Chaetoceros decipiens | 0.92 | 0.78 | 0.85 | 0.0011 | ** |
| 3-20 $\mu\text{m}$ | MIZ+OW | asv_023_00225 | Pelagophyceae | Ankylochrysis sp. | 0.96 | 0.53 | 0.72 | 0.0116 | * |
| 3-20 $\mu\text{m}$ | MIZ+OW | asv_023_00283 | Bacillariophyta | Attheya septentrionalis | 0.92 | 0.34 | 0.56 | 0.0476 | * |
| 3-20 $\mu\text{m}$ | MIZ+OW | asv_023_00218 | Bacillariophyta | Chaetoceros danicus | 0.92 | 0.69 | 0.8 | 0.0028 | ** |
| 3-20 $\mu\text{m}$ | MIZ+OW | asv_023_00223 | Prymnesiophyceae | Prymnesiophyceae Clade F XX sp. | 0.9 | 0.5 | 0.67 | 0.0302 | * |
| 3-20 $\mu\text{m}$ | MIZ+OW | asv_023_00013 | Bacillariophyta | Thalassiosira rotula | 0.97 | 0.31 | 0.55 | 0.0398 | * |
| > 20 $\mu\text{m}$ | UI | asv_023_00086 | Prymnesiophyceae | Chrysochromulina sp. | 0.96 | 0.17 | 0.41 | 0.017 | * |
| > 20 $\mu\text{m}$ | UI | asv_023_00046 | Bacillariophyta | Pseudo-nitzschia seriata | 0.79 | 0.85 | 0.82 | 1e-04 | *** |
| > 20 $\mu\text{m}$ | UI | asv_023_00105 | Prymnesiophyceae | Phaeocystis cordata | 0.76 | 0.45 | 0.58 | 0.0019 | ** |
| > 20 $\mu\text{m}$ | MIZ | asv_023_00259 | Bacillariophyta | Entomoneis ornata | 0.83 | 0.62 | 0.72 | 2e-04 | *** |
| > 20 $\mu\text{m}$ | MIZ | asv_023_00265 | Bacillariophyta | Pauliella toeniata | 0.69 | 0.68 | 0.68 | 0.0046 | ** |
| > 20 $\mu\text{m}$ | OW | asv_023_00334 | Bacillariophyta | Chaetoceros contortus | 0.72 | 0.83 | 0.77 | 1e-04 | *** |
| > 20 $\mu\text{m}$ | OW | asv_023_00407 | Bacillariophyta | Chaetoceros diadema 1 | 0.82 | 0.67 | 0.74 | 1e-04 | *** |
| > 20 $\mu\text{m}$ | MIZ+UI | asv_023_00009 | Bacillariophyta | Melosira arctica | 1 | 1 | 1 | 1e-04 | *** |
| > 20 $\mu\text{m}$ | MIZ+UI | asv_023_00025 | Bacillariophyta | Melosira arctica | 1 | 0.81 | 0.9 | 1e-04 | *** |
| > 20 $\mu\text{m}$ | MIZ+UI | asv_023_00038 | Cryptophyceae | Teleaulax gracilis | 0.97 | 0.51 | 0.71 | 9e-04 | *** |
| > 20 $\mu\text{m}$ | MIZ+UI | asv_023_00049 | Bacillariophyta | Navicula sp. | 0.97 | 0.84 | 0.9 | 1e-04 | *** |
| > 20 $\mu\text{m}$ | MIZ+UI | asv_023_00064 | Bacillariophyta | Fragilaria sp. | 0.98 | 0.73 | 0.84 | 1e-04 | *** |
| > 20 $\mu\text{m}$ | MIZ+UI | asv_023_00073 | Bolidophyceae | Triparma laevis clade | 0.99 | 0.61 | 0.78 | 1e-04 | *** |
| > 20 $\mu\text{m}$ | MIZ+UI | asv_023_00075 | Dictyochophyceae | Dictyocha speculum | 1 | 0.8 | 0.89 | 1e-04 | *** |
| > 20 $\mu\text{m}$ | MIZ+UI | asv_023_00084 | Bacillariophyta | Raphid-pennate X sp. | 1 | 0.81 | 0.9 | 1e-04 | *** |
| > 20 $\mu\text{m}$ | MIZ+UI | asv_023_00099 | Bacillariophyta | Cylindrotheca closterium | 1 | 0.57 | 0.75 | 1e-04 | *** |
| > 20 $\mu\text{m}$ | MIZ+UI | asv_023_00178 | Bacillariophyta | Raphid-pennate X sp. | 1 | 0.31 | 0.56 | 0.0124 | * |
| > 20 $\mu\text{m}$ | MIZ+UI | asv_023_00172 | Bacillariophyta | Attheya septentrionalis | 1 | 0.8 | 0.89 | 1e-04 | *** |
| > 20 $\mu\text{m}$ | MIZ+UI | asv_023_00328 | Bacillariophyta | Nitzschia sp. | 1 | 0.34 | 0.58 | 0.0166 | * |

**Table 4:** (continued)
| Size fraction | Sectors | ASVs | Class | Species | A | B | Stat | p.value | Sign |
| --- | --- | --- | --- | --- | --- | --- | --- | --- | --- |
| > 20 $\mu\text{m}$ | MIZ+UI | asv_023_00184 | Bacillariophyta | Pleurosigma intermedium | 1 | 0.65 | 0.8 | 1e-04 | *** |
| > 20 $\mu\text{m}$ | MIZ+UI | asv_023_00219 | Bacillariophyta | Bacillaria paxillifer | 1 | 0.5 | 0.71 | 3e-04 | *** |
| > 20 $\mu\text{m}$ | MIZ+UI | asv_023_00268 | Bacillariophyta | Stauroneis kriegeri | 1 | 0.49 | 0.7 | 0.001 | *** |
| > 20 $\mu\text{m}$ | MIZ+UI | asv_023_00277 | Bacillariophyta | Naviculales sp. | 0.97 | 0.61 | 0.77 | 3e-04 | *** |
| > 20 $\mu\text{m}$ | MIZ+UI | asv_023_00308 | Bacillariophyta | Synedra hyperborea | 1 | 0.42 | 0.65 | 0.0033 | ** |
| > 20 $\mu\text{m}$ | MIZ+UI | asv_023_00335 | Bacillariophyta | Raphid-pennate X sp. | 1 | 0.61 | 0.78 | 1e-04 | *** |
| > 20 $\mu\text{m}$ | MIZ+OW | asv_023_00001 | Prymnesiophyceae | Phaeocystis pouchetii | 0.91 | 1 | 0.96 | 1e-04 | *** |
| > 20 $\mu\text{m}$ | MIZ+OW | asv_023_00013 | Bacillariophyta | Thalassiosira rotula | 0.96 | 0.96 | 0.96 | 1e-04 | *** |
| > 20 $\mu\text{m}$ | MIZ+OW | asv_023_00028 | Bacillariophyta | Thalassiosira sp. | 0.77 | 0.85 | 0.81 | 3e-04 | *** |
| > 20 $\mu\text{m}$ | MIZ+OW | asv_023_00057 | Bacillariophyta | Thalassiosira sp. | 0.93 | 0.9 | 0.92 | 1e-04 | *** |
| > 20 $\mu\text{m}$ | MIZ+OW | asv_023_00071 | Bacillariophyta | Thalassiosira anguste-lineata | 0.97 | 0.71 | 0.83 | 1e-04 | *** |
| > 20 $\mu\text{m}$ | MIZ+OW | asv_023_00079 | Bacillariophyta | Eucampia sp. | 0.94 | 0.9 | 0.92 | 1e-04 | *** |
| > 20 $\mu\text{m}$ | MIZ+OW | asv_023_00111 | Bacillariophyta | Chaetoceros cinctus | 0.89 | 0.81 | 0.85 | 1e-04 | *** |
| > 20 $\mu\text{m}$ | MIZ+OW | asv_023_00136 | Bacillariophyta | Chaetoceros decipiens | 0.87 | 0.79 | 0.83 | 1e-04 | *** |
| > 20 $\mu\text{m}$ | MIZ+OW | asv_023_00137 | Bacillariophyta | Detonula confervacea | 0.9 | 0.79 | 0.84 | 1e-04 | *** |
| > 20 $\mu\text{m}$ | MIZ+OW | asv_023_00177 | Bacillariophyta | Chaetoceros rostratus | 0.89 | 0.85 | 0.87 | 1e-04 | *** |
| > 20 $\mu\text{m}$ | MIZ+OW | asv_023_00218 | Bacillariophyta | Chaetoceros danicus | 0.92 | 0.73 | 0.82 | 1e-04 | *** |

**3-20** *µ***m**. There was no significant association between taxa and the UI sector in the 3-20 *µ*m size fraction, and the six ASVs that were considered indicators of this sector presented lower *p*-values, with *Bacillaria paxillifer* (*p*-value = 0.003) and *Pterosperma* sp. (*p*-value = 0.0095) presenting the highest association score (Table 4). Considering only highly significant associations (*p*-value < 0.001), *Navicula* sp. (ASV_0049) was the only ASV representative of the MIZ, while several Ochrophyta and one Cryptophyta member were identified as indicators from the MIZ+UI sector group, all of them highly significant related to ice-associated sectors also in the 0.2-3 *µ*m size fraction: *F. daucoides* (ASV_0041), *Fragilariopsis cylindrus* (ASV_0015), *Pseudonitzschia seriata* (ASV_0046), *C. neogracilis* (ASV_0048), MOCH-2 sp. (ASV_0061), *Triparma laevis* clade (ASV_0073) and *Dictyocha speculum* (ASV_0075). Two centric diatoms were significantly associated with the OW sector, *Thalassiosira* sp. (ASV_0057) and *Chaetoceros rostratus* (ASV_0177), and another centric diatom to the MIZ+OW sector group, *Eucampia* sp. (ASV_0079).

**> 20** *µ***m**. From the 30 highly significant indicator ASVs found in the > 20 *µ*m size fraction, only one was related to the UI (*Pseudo-nitzschia seriata*, ASV_0046), one to the MIZ (*Entomoneis ornata*, ASV_0259), and two to the OW sector (*Chaetoceros contortus* ASV_0334 and *Chaetoceros diadema* 1 ASV_0407) (Table 4). Fifteen ASVs were highly related to the MIZ+UI sector group, including two *M. arctica* ASVs (0009 and 0025) and some also related to ice-associated sectors in the other size fractions: *Navicula* sp. (ASV_0049), *Triparma laevis* clade (ASV_0073), *Dictyocha speculum* (ASV_0075), and *Bacillaria paxillifer* (ASV_0219). From the eleven indicator ASVs strongly associated with the MIZ+OW sector group, ten were centric diatoms, including four *Chaetoceros*, four *Thalassiosira*, one *Eucampia* sp. and one *Detonula confervacea* (ASV_0137). Interestingly, the most abundant ASV from the whole dataset, *P. pouchetii* (ASV_0001), was also highly related to the MIZ+OW sector group in the > 20 *µ*m size fraction (Table 4).

### Distribution of abundant ASVs

The distribution of the ten most abundant ASVs within each division demonstrated that although the dominant community might be comprised of few genera, ASV-level distribution follows distinct patterns within these genera and even within the same species (Figure 6). It is also notable that some ASVs were particularly adapted to distinct environments, regardless of differences between sectors, since they were present (and abundant) throughout the dataset, such as *M. polaris* (ASV_0003), *Teleaulax glacialis* (ASV_0038) and *P. pouchetii* (ASV_0001) (Figure 6). Chlorophyta top 10 ASVs belonged to five genera: *Bathycoccus*, *Mantoniella*, *Micromonas*, *Pterosperma* and *Pyramimonas*, from which five ASVs were significantly correlated to ice-associated sectors. *M. polaris* ASV_0154 had a single base pair difference with *M. polaris* ASV_0003 (Figure S6) and was less abundant than the latter in our dataset (Figure 6), as well as in other Arctic datasets (Figure S7). *M. squamata* (ASV_0104) and *Pterosperma* sp. (ASV_0244) were present in UI samples, mainly within 0.2-3, but also in the 3-20 *µ*m size fraction.

**Figure 6:**
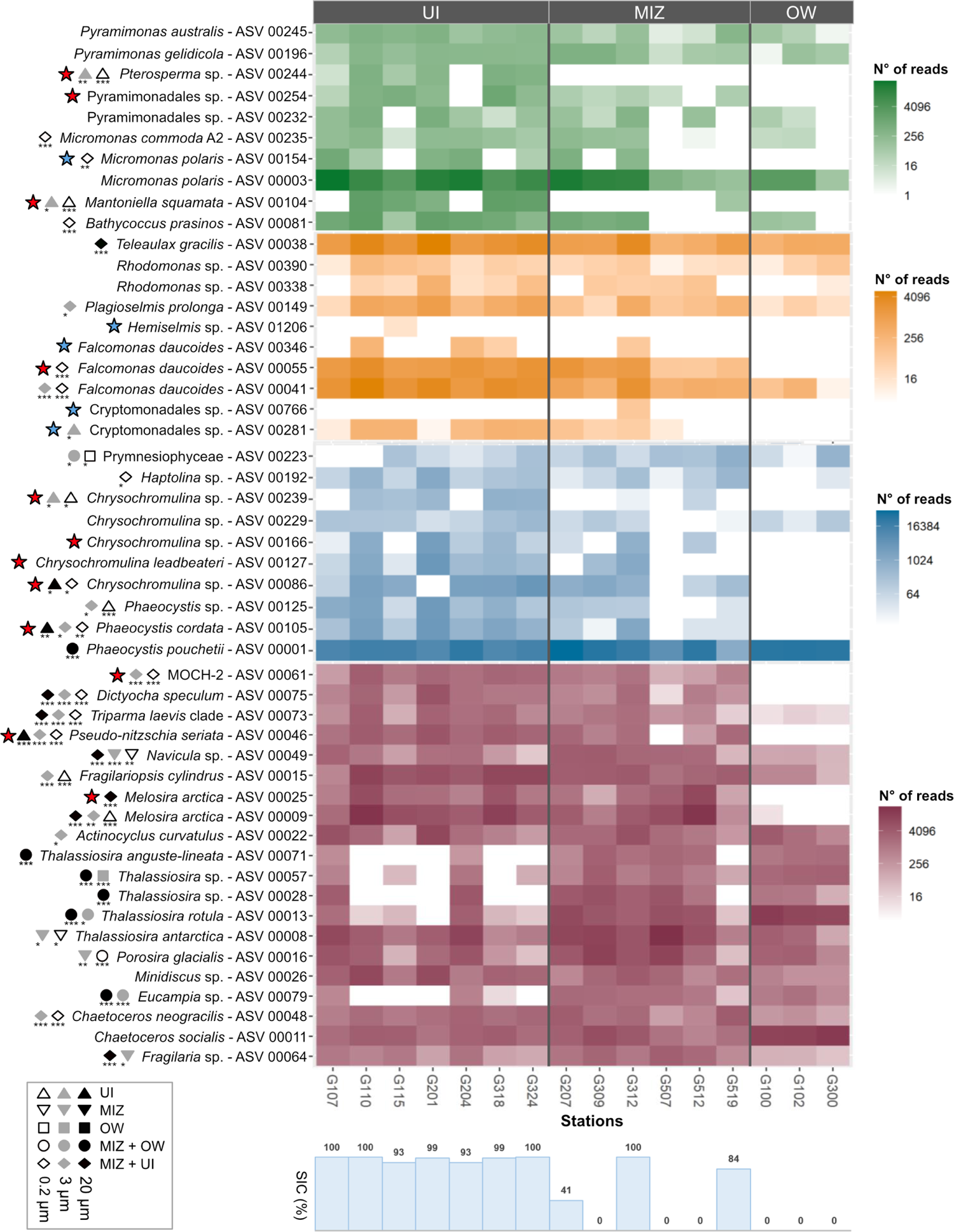
Taxa distribution of the most abundant ASVs for each sector, divided by sampling stations; the top 10 ASVs were selected within Chlorophyta (green), Cryptophyta (orange), Haptophyta (blue), and the top 20 most abundant within the highly diverse Ochrophyta division (red); symbols indicate if a given ASV was reported as indicator ASV for UI (triangles), MIZ (inverted triangles), OW (squares), MIZ+UI (diamonds) or MIZ+OW (circles) sector groups within 0.2-3 *µ*m (white), 3-20 *µ*m (grey) or > 20 *µ*m (black) size fractions; asterisks indicate the *p*-values associated with the indicator ASV: 0 (***), 0.001 (**), and 0.01 (*); red and blue stars indicate if a given ASV was found exclusively in ice-associated sectors, being blue stars not abundant ASVs; SIC = sea-ice concentration on each sampling station.

Within the top 10 most abundant Cryptophyta ASVs, three were assigned as *F. daucoides*. While ASV_0041 *F. daucoides* was strongly associated with MIZ+UI samples for both 0.2-3 and 3-20 *µ*m size fractions but also found in the OW sector, ASV_0055 was exclusively found in ice-associated sectors in the smaller size fraction (Figure 6). *T. gracilis* ASV_0038 was highly abundant at all sectors, but was flagged as indicator ASV for the MIZ+UI sector group in the > 20 *µ*m size fraction.

Five *Chrysochromulina* and three *Phaeocystis* ASVs comprised the ten most abundant Haptophyta, three of them only found in ice-associated sectors. Interestingly, *P. cordata* ASV_0105 and *Phaeocystis* sp. ASV_0125 were strongly associated with both UI or MIZ+UI sector group, while *P. pouchetii* ASV_0001 was associated with the larger size fraction of MIZ+OW, although highly abundant at all stations (Figure 6).

The most abundant non-diatom Ochrophyta were MOCH-2 ASV_0061, *D. speculum* ASV_0075 and *T. laevis* ASV_0073, all of them flagged as ice-associated indicator ASVs in all size fractions, except for MOCH-2, which was not an indicator for the > 20 *µ*m size fraction (Figure 6). Many diatoms were flagged as indicator species for ice-associated sectors, including pennate diatoms such as *P. seriata*, *Navicula* sp., and *Fragilaria* sp., and centric diatoms such as *M. arctica* and *C. neogracilis*. Interestingly, from the five *Thalassiosira* ASVs ranking as most abundant Ochrophyta, four were considered indicator species of OW or MIZ+OW sectors, while *Thalassiosira antarctica* was mainly associated with the smaller size fractions of the MIZ sector, although present at all sectors. Other centric diatoms were also indicators of the MIZ+OW sectors, such as *Eucampia* sp. ASV_0079 and *P. glacialis* ASV_0016 ASV_0073 (Figure 6). The distribution of three abundant *Thalassiosira* flagged as OW or MIZ+OW indicators ASVs have been previously described as having a broad distribution in both lower and higher latitudes in relation to the sampled area from the present study (Figure S8).

## Discussion

### General bloom progression

Under-ice blooms are reported from throughout the Arctic, but the onset conditions, biogeochemical dynamics and taxa succession are subjected to regional features (Ardyna et al. 2020). Although the Green Edge cruise transects were east-west spatial snapshots, the sampling strategy recovered nutrients, cell abundance, plankton diversity and metabolism variability as the bloom progressed from UI to OW (Lafond et al. 2019; Randelhoff et al. 2019; Saint-Béat et al. 2020; Vilgrain et al. 2021). During pre-bloom conditions, photosynthetic activity is limited by light availability, and under-ice populations are shade-acclimated (Ardyna et al. 2020). During the Green Edge campaign, however, light availability and vertical mixing should have permitted a bloom initiation under nearly 100% sea-ice cover (UI sector), but the bloom peaked in terms of chlorophyll-a approximately 10 days after ice retreat (Randelhoff et al. 2019), around the limit between the MIZ and the OW sector.

In general, FCM abundance data indicated higher pico- and nano-phytoplankton abundances within the MIZ and the OW sectors, respectively. Although the micro-phytoplankton size fraction was not counted by FCM, the amplicon data suggests a community structure shift from smaller to larger size fractions as the bloom progresses. FCM data shows that while UI pico-phytoplankton reached its maximum and was relatively well distributed down the euphotic zone of the water column, nano-phytoplankton community was particularly abundant right beneath the sea-ice, indicating a close association between the top of the water column and the bottom of the sea-ice (Figure 3). Interestingly, contrary to cell abundance, the relative abundance of the main genera did not seem to change with depth within the UI sector, except for a higher contribution of *Phaeocystis* spp. and *Teleaulax* sp. reads in surface. Diversity decreased eastward from the UI to the OW sector, which represents the different stages of the phytoplankton spring bloom but is also influenced by the different water masses within Baffin Bay. Ice-associated stations, most still covered with sea-ice and categorized as low-productivity stations by Lafond et al. (2019), harbored the most diverse community, from the genus to the ASV level, and within every size fraction (Figure 4, Table 3). Under-ice communities are adapted to low-light environments, capable of maximizing light absorption by increasing intracellular concentrations of accessory and photosynthetic pigments (Lewis et al. 2019). Smaller phytoplankton cells from the Beaufort Sea were reported to be more efficient at harvesting light due partially to an increase in chlorophyll *b* content, which is associated with the low light conditions in autumn and winter (Matsuoka et al. 2009). Such recruitment due to polar dark conditions might explain the dominance of *Micromonas*, an early-bloom taxa (Lovejoy et al. 2007) with known persistence during winter (Joli et al. 2017; Vader et al. 2015), within the UI and MIZ sectors, which represented almost the total bulk of pico-sized plankton. Several abundant non-diatom ASVs were flagged as indicator ASVs from ice-associated sectors, especially in the smaller size fraction (Figure 6). Many of these indicator ASVs were found in datasets from the high Arctic (see ASV diversity subsection below), suggesting that UI and MIZ phytoplankton communities are highly diverse, probably low-light adapted populations of smaller organisms, which seem to be connected to higher-latitude communities (Kalenitchenko et al. 2019) probably via water mass intrusions from the Nares Strait and the Smith, Jones and Lancaster Sounds (Bluhm et al. 2015; Tang et al. 2004).

We observed a general increase in pico- and nano-phytoplankton cells within MIZ, with a sharp decline towards the OW sector. The MIZ sector showed evidence of increased biological activity: pico- and nano-phytoplankton abundance, dissolved and particulate organic carbon concentration, particulate organic nitrogen concentration, dissolved organic nitrogen release and primary production in general were higher in the MIZ than in the UI or OW sectors. Vilgrain et al. (2021) reported that near the ice-edge, copepods were heavily pigmented due partially to full gut content. In the present study, the relatively steady community composition between UI and MIZ, which comprised the peak of the bloom, may be explained by the seeding of taxa through ice melt water (Mundy et al. 2011) combined with the “priming” effect suggested by Lewis et al. (2019). The “priming” effect arises from the acclimation of pre-bloom, under-ice communities to low and highly variable light input due to patchy snow cover and melt-pond/open water leads formation, resulting in a competitive advantage to rapidly exploit increasing irradiation (Lewis et al. 2019).

As the phytoplankton spring bloom progresses, the phytoplankton community must transition from light-limited conditions, characteristic of a pre-bloom state, to a high-light, nutrient-limited environment (Lewis et al. 2019). The nitracline deepened from 0 in UI to more than 30 meters in OW, along with the development of a subsurface chlorophyll *a* maximum (Randelhoff et al. 2019), as a result of the rapid consumption of inorganic nutrients in surface, following an expected trend in Arctic plankton phenology (Ardyna et al. 2020; Martin et al. 2010). In the post-bloom conditions such as found within the OW sector, new production is confined to deeper layers, and the euphotic layer is then dominated by regenerative production (Sakshaug 2004). Within ice-free Arctic waters, species might also be subjected to detrimental effects of high UV exposure resulting in low cell viability and photosynthetic performance decline, a scenario where, in general, diatoms out-compete other microalgae (Alou-Font et al. 2016). The increase in *Chaetoceros* spp. observed here in the OW sector, especially in the upper layers within the 3-20 *µ*m size fraction, may reflect an ecological advantage within a post-bloom scenario, since this genus has a high growth irrespective of nitrogen source in polar (Schiffrine et al. 2020) and subtropical (Morando and Capone 2018) environments. The only abundant diatoms flagged as indicator ASVs for OW or MIZ+OW sectors belonged to *Thalassiosira* and *Eucampia*, two genera that were also reported to be effective scavengers for different nitrogen sources, out-competing other plankton community members within nutrient poor waters (Morando and Capone 2018). NMDS analysis highlighted the gradient from the UI sector with higher concentration of major nutrients such as nitrate and phosphate, to the MIZ/OW sector with higher ammonium and urea assimilation, together with an increase of large (> 20 *µ*m) diatoms. Besides the bottom-up pressure favoring such specialized taxa, the community structure responsible for the secondary production within the OW sector might also play a role in the differential top-down control of smaller taxa, since, in the westernmost stations, the copepod populations were dominated by smaller organisms, in particular nauplii and young copepod stages (Vilgrain et al. 2021).

### Autotroph microdiversity

Different ASVs within the same species can represent distinct ecotypes, leading to resilience and adaptation of microbial populations under changing environmental conditions (García-García et al. 2019; Needham and Fuhrman 2016), and the persistence of particular lineages over time. We have identified several species comprising more than one ASV. Sjöqvist & Kremp (2016) reported that genetic diversity within diatom species ensured an optimized ecological performance, including carbon uptake and overall resistance to environmental changes.

### Diatoms

Sea-ice has been long recognized as an important substrate for marine diatoms (Horner et al. 1992; Poulin et al. 2011), where brine channels and pockets serve as habitat for a specific interstitial and sub-ice community. The centric diatom *M. arctica* is an ice-associated taxon, forming long strands in the water column attached to the sea-ice (Poulin et al. 2014; Wassmann et al. 2006). This diatom is readily released as ice melts, rapidly sinking and forming vast sea-floor deposits down to more than 4,000 m depth, with a high impact on carbon export and benthic fauna (Boetius et al. 2013). Our data indicated two highly abundant *M. arctica* ASVs not only in UI but also in the MIZ sector, the latter with several ice-free stations. Sampling roughly the same stations and using microscopy and pigment-based analysis, Lafond et al. (2019) showed that *M. arctica* in the MIZ was mostly in the form of actively silicifying resting spores, reaching up to 82% of biogenic silica production. The high degree of significance associating *M. arctica* with ice-sectors in the present study corroborate its assignation as a sea-ice specialist taxa, but the high number of reads in the intermittently ice-covered Baffin Bay challenges previous work hypothesizing its preference for multi-year sea-ice (Hop et al. 2020).

Lafond et al. (2019) observed that diatoms during the melting season in Baffin Bay formed two distinct community clusters: one less productive, associated with Pacific-originating waters, and another associated with the core of the diatom bloom, within Atlantic-influenced stations, mainly in open waters or at an advanced state of sea-ice melting. Our data are consistent with the current view of the succession pattern of the Arctic diatom bloom, where ice-associated early bloom stages are characterized by a higher diversity of pennate diatoms while its full development consists of larger centric diatoms, such as *Thalassiosira* and *Chaetoceros* (Oziel et al. 2019). Such dynamics are partly explained by differences in nutrient acquisition strategies (Morando and Capone 2018) and by increased photochemical damage experienced by sympagic pennate diatoms under ice-free, high-luminosity environments (Kvernvik et al. 2020). In the present study, abundant pennate diatoms, *P. seriata* ASV_0046, *Navicula* sp. ASV_0049, and *Fragilaria* sp ASV_0064, were all flagged as indicator species for ice-associated sectors. Interestingly, one of the most abundant *Chaetoceros* ASVs (*C. neogracilis* ASV_0048) was considered a highly significant ice-associated indicator ASV, but only for the smaller size fraction. *C. neogracilis* is a species complex with at least four known clades which share identical 18S rRNA sequences (Balzano et al. 2017), so it is possible that the *C. neogracilis* distribution observed in the present study is masking a finer clade-specific distribution.

One of the hypothesis from the present work was that taxa significantly associated with Atlantic-influenced east Baffin Bay could be due to Atlantification processes that include input of warm-adapted taxa from the eastern side of Davis Strait, which could thrive after ice melt. However, OW+MIZ indicator taxa that made up the abundant community, such as *Thalassiosira anguste-lineata* ASV_0071, *Thalassiosira* sp. ASV_0057, and *T. rotula* ASV_0013, although sometimes displaying a wider distribution towards lower latitudes, are considered part of the Arctic phytoplankton community, as shown by their distribution over many polar studies in metaPR^2^ (Figure S8). Those ASVs were also found within UI sector, indicating that their attribution as MIZ+OW indicator taxa was mainly due to a better adaptation to low nutrient post-bloom conditions.

### Non-diatom Ochrophyta

Although Lafond et al. (2019) identified the core of the diatom bloom within Atlantic-influenced Baffin Bay, our FCM and metabarcoding data indicate the importance of smaller size fractions within ice-associated sectors in terms of cell abundance and overall plankton diversity. For example, nano-phytoplankton Ochrophyta diversity within OW was dominated by a single diatom genus, *Chaetoceros*, while in the UI and MIZ both diatom and non-diatom Ochrophyta were much more diverse, with high abundances of MOCH-2, *Dictyocha speculum*, *Triparma laevis* clade, flagged as indicator ASVs for ice-associated sectors along with an unidentified Florenciellales.

The presence of the silicoflagellate *D. speculum* (synonym of *Octactis speculum*, Chang et al. 2017) in Arctic waters was first reported in the region more than a century ago (Lovejoy et al. 2002) and since regularly cited in the literature (Crawford et al. 2018). Its assignation as an indicator ASV for ice-associated sectors within all size classes in the present study might be a consequence of the presence of several life stages, including amoeboid, multinucleate and skeleton-bearing stages with different cell sizes (Chang et al. 2017; Moestrup and Thomsen 1990). Studying a 35-year sampling series, Hop et al. (2020) reported that this species has a higher frequency of occurrence in multiyear ice (24 %) in comparison to first-year ice samples (6 %).

### Chlorophyta

The most striking difference between ASV distribution patterns among a dominant species in the present study was observed within *M. polaris* populations. The Chlorophyta genus *Micromonas* is diverse and widely distributed from coastal to oceanic waters through all the global latitudinal ranges (Simon et al. 2017; Tragin and Vaulot 2019). It exhibits a wide thermal niche and is considered a sentinel for polar (Freyria et al. 2021) and global plankton diversity (Demory et al. 2019) in relation to temperature changes in the oceans. The use of metabarcoding datasets combined with microdiversity approaches has previously allowed the discovery of new polar *Micromonas* ecotypes, such as the *Micromonas* B3 clade, which displays a wider distribution band towards lower latitudes than *M. polaris* (Tragin and Vaulot 2019).

The *M. polaris* CCMP2099 strain, isolated from North Water Polynya (Lovejoy et al. 2007) and the RCC2306 strain from the Beaufort Sea (holotype of the species, Simon et al. 2017) are 100% similar in the V4 region of the 18S rRNA to the *M. polaris* ASV_0003 from the present study. *M. polaris* ASV_0003 has a widespread distribution pattern with a high abundance in all sectors, in accordance with its dominant role within the Arctic (Balzano et al. 2012; Lovejoy et al. 2002, 2007; Not et al. 2005). Although *M. polaris* ASV_0154 differs from ASV_0003 by a single nucleotide (Fig. S6), its significantly different distribution (Fig. S7), and the assignation of ASV_0003 as an indicator species, suggests it might represent a new ecotype. There is no 100% similarity match in GenBank with *M. polaris* ASV_0154, either to strains or environmental sequences. The distribution of *M. polaris* ASV_0154 is pan-Arctic, although it always contributes to a small fraction of *Micromonas* reads. *M. polaris* ASV_0154 has also been found in the Nares Strait (metaPR^2^ set #42, Kalenitchenko et al. 2019) (Figure S7), also comprising a small fraction of *Micromonas* reads. The Nares Strait is connected to northern Baffin Bay and is responsible for southward transport of waters and ice from the Arctic Ocean into the region (Tang et al. 2004). Using a decade-long 18S rRNA data series, Freyria et al. (2021) have identified *M. polaris* as a summer specialist favored by nutrient-poor waters, in contrast to the present study, which identifies the species either as a generalist, present in all sectors (ASV_0003) or as an ice-associated indicator ASV (ASV_0154). The difference in *M. polaris* distribution patterns between Freyria et al. (2021) and the present study might be related to the distinct geographic location, time of sampling and data processing. Freyria et al. (2021) sampled the northernmost sector of Baffin Bay, more precisely the North Water, a hydrographically distinct region, close to 77*^◦^* N, bordered by Ellesmere Island and Greenland. The authors sampled the transition from summer to autumn, while the present study focus on the transition between spring and summer. *M. polaris* abundance was previously reported decreasing in numbers and activity towards winter, partially due to vulnerability to specific viral infection during this period, and then recovering rapidly even at low irradiances (Joli et al. 2017).

One Chlorophyta ASV was associated with under-ice samples: *Pterosperma* sp. ASV_0244, which was found exclusively in the UI sector, and flagged as one of the few abundant indicator species from it, and not from the MIZ+UI sector group. Although the genus *Pterosperma* has been reported from several regions within the Arctic (Joli et al. 2017; Lovejoy et al. 2002), with a preference for multi-year ice over first-year ice (Hop et al. 2020), there was no 100% match between ASV_0244 and any strain or environmental sequence in GenBank. *Pterosperma* sp. ASV_0244 was only found in three other samples from the 41 global datasets in the metaPR^2^ database, in the east coast of Greenland (Kopf et al. 2015) and the Nansen Basin (Metfies et al. 2016). Although previous studies have identified sea-ice associated communities harboring a high relative abundance of *Pyramimonas* (Gradinger 1996; Mundy et al. 2011), our indicator analysis did not detect any specific distribution linked to sea ice for this genus.

### Haptophyta

*P. pouchetii* is ubiquitous throughout the Arctic (Lasternas and Agustı 2010; Schoemann et al. 2005) and has been reported to be capable of early blooms, even under snow-covered ice pack (Assmy et al. 2017). Although *P. pouchetii* ASV_0001 reads were present in all size fractions and sectors, the species was flagged as an indicator ASV for the MIZ+OW sector in the > 20 *µ*m size fraction. This might indicate a prevalence of large *P. pouchetii* colonies towards eastern side of Baffin Bay, as the bloom progresses. In general, blooming species of the genus *Phaeocystis* increase their C:N ratios under high light/low nutrient conditions, mainly through the production of the polysaccharide-based mucilaginous matrix embedding colonies reaching up to 3 cm, which serve as energy storage and a defense against grazers (Schoemann et al. 2005) and references therein). The dominance of *P. pouchetii* colonial form was reported during the Arctic 2007 ice-melt record (Lasternas and Agustı 2010), while the single-cell form was reported during overwintering (Vader et al. 2015). The fact that *P. pouchetii* is adapted to grow in nutrient-replete waters with 100% sea-ice cover such as found in the UI sector to the nutrient-depleted/high light OW sector corroborates earlier studies identifying this taxa as a potential winner for future Arctic scenarios, where its plasticity regarding life cycle stages, and flexibility for light and nutrient uptake, as well as resistance to zooplankton grazing will likely impact polar phytoplankton community structuring, trophic energy transfer and carbon export (Lasternas and Agustı 2010; Verity et al. 2007; Wassmann et al. 2006).

The higher relative contribution of *Chrysochromulina* in deeper samples agrees with previous reports linking this genus with deep chlorophyll maximum communities (Balzano et al. 2012). Many *Chrysochromulina* spp. ASVs were exclusively found or flagged as indicator taxa for ice-associated sectors. *Chrysochromulina* frequently occurs in sympagic communities and is considered one of the few ice-associated haptophytes (Mundy et al. 2011), but it is also present in ice-free waters from the Arctic (Balzano et al. 2012; Lovejoy et al. 2002) and the Antarctic (Luo et al. 2016; Trefault et al. 2021). The fact that the genus *Chrysochromulina* is morphometrically highly diverse (Egge et al. 2014) down to the subspecies level (Balzano et al. 2012; Needham and Fuhrman 2016) implies that monitoring diversity should comprise high-resolution analysis of species/ecotypes distributions. Although not really abundant in the present study, *Chrysochromulina* sp. ASV_0542 was found exclusively in under-ice samples. It was found previously to reach up to 2% of total eukaryotic reads in the Nares Strait (metaPR^2^ set #42, Kalenitchenko et al. 2019), but not detected elsewhere.

The distribution of *Pterosperma* sp. ASV_0244 and *Chrysochromulina* sp. ASV_0542 restricted to a limited Arctic latitudinal band suggests a narrow ecological niche, and its significant association with ice-covered sites from the present study indicate that those taxa might be good proxies for diversity changes within the region. Some taxa flagged as indicator ASVs for ice-associated sectors do have a broader range of distribution towards lower latitudes, such as *Phaeocystis* ASV_0125 and *M. commoda* A2 ASV_0235. Whether this distribution results from the combination of different ecotypes masked by the resolution of the marker gene, or simply represents taxa out-competed within an ice-free, high-light and low nutrient environment, is still an open question.

## Conclusions

Taking the spatial transects as a temporal snapshot of the phytoplankton spring bloom dynamics over Baffin Bay, we observed a shift from a highly diverse under-ice and ice-edge community comprised of smaller taxa, to a low-diversity, highly specialized community where larger centric diatoms and *P. pouchetii* were better adapted to the harsh high light/low nutrient post-bloom environment. The taxa abundant in the ice-free Atlantic-influenced Baffin Bay were in general also present in Arctic-influenced sectors and other polar studies, indicating that the advection of warm-adapted taxa by Atlantic inflow was not detectable or significant in this region. This conclusion must be taken with caution, however, due to the limited space and time sampling range of the present study. The presence of taxa with intra-species variability such as *F. daucoides*, *M. arctica* and *M. polaris* reinforces the urgency of renewed culturing efforts to better comprehend its ecological impacts. Although thinner sea-ice might increase the magnitude of the sub-ice blooms of taxa with a high carbon export rate such as *M. arctica* (Poulin et al. 2014) earlier in the season, our data indicate that as Baffin Bay ice cover shrinks sooner and faster with spring blooms onset (Stroeve et al. 2014), this might lead to widespread post-bloom conditions dominated by a much less diverse community, which might have implications in the recruitment of sympagic communities in subsequent years.

## Acknowledgements

We wish to thank officers and crew of CCGS Amundsen. The success of the field campaign was widely contributed to G. Bécu, J. Lagunas, D. Christiansen-Stowe, J. Sansoulet, E. Rehm, M. Benoît-Gagné, M.-H. Forget and F. Bruyant from Takuvik laboratory and J. Bourdon, C. Marec and M. Picheral from CNRS. We also thank Québec-Océan and the Polar Continental Shelf Program for their in-kind contribution in terms of polar logistics and scientific equipment. We thank the ABIMS platform of the FR2424 (CNRS, Sorbonne Université) for bioinformatics resources. We thank Margot Tragin for her help during sampling and onboard sample processing.

## Author contributions statement

CGR, ALS, DM and DV processed the samples and produced data; CGR, DV, NT, CL and ALS analyzed and interpreted data; CGR and DV wrote the paper; All authors read and approved the final manuscript.

## Funding information

Catherine Gérikas Ribeiro was supported by the FONDECYT Project No. 3190827. The Green Edge project was funded by the following French and Canadian programs and agencies: ANR (Contract #111112), ArcticNet, CERC on Remote sensing of Canada’s new Arctic frontier, CNES (project #131425), French Arctic Initiative, Fondation Total, CSA, LEFE and IPEV (project #1164). This project was conducted using the Canadian research icebreaker CCGS Amundsen with the support of the Amundsen Science program funded by the Canada Foundation for Innovation (CFI) Major Science Initiatives (MSI) Fund. The project was conducted under the scientific coordination of the Canada Excellence Research Chair on Remote sensing of Canada’s new Arctic frontier and the CNRS & Université Laval Takuvik Joint International Laboratory (UMI3376). Nicole Trefault was supported by Fondecyt No. 1190879 and INACH RT_34-17.

## Additional information

To include, in this order:

**Accession codes** (where applicable);

## Data availability

Physicochemical and biological data from the Green Edge project are available at http://www.obs-vlfr.fr/proof/php/GREENEDGE/x_datalist_1.php?xxop=greenedge&xxcamp=amundsen and at https://www.seanoe.org/data/00487/59892/ as described in detail in Bruyant et al. (2022). Raw metabarcoding sequences are available under the Accession Code PRJNA810033 from GenBank SRA. Source code for sequence processing is available at https://github.com/vaulot/Paper-2021-Vaulot-metapr2/tree/main/R_processing.

## Competing interests

The authors declare no competing financial interests.

## ORCID Numbers

- Catherine Gérikas Ribeiro: 0000-0003-0531-2313
- Daniel Vaulot: 0000-0002-0717-5685
- Nicole Trefault: 0000-0002-4388-6791
- Connie Lovejoy: 0000-0001-8027-2281
- Adriana Lopes dos Santos: 0000-0002-0736-4937

## Supplementary material

All supplementary material is available at https://github.com/catherine-gerikas/GE_Amundsen_18S_metaB_supplementary_material

### Supplementary Data

Supplementary Data S1: Sample dates and environmental data available. Available in: https://github.com/catherine-gerikas/GE_Amundsen_18S_metaB_supplementary_material

### Supplementary Tables

**Table S1:** List of variables measured during the Green Edge cruise (see Data Set S1).

| Variable | Description | Unit |
| --- | --- | --- |
| sample_code | sample code |  |
| fraction_name | size fraction |  |
| station_id | station ID |  |
| CTD | ID of CTD cast |  |
| transect | cruise transect ID |  |
| bot_depth | bottom depth at a given station | m |
| depth | depth on each the sample was taken | m |
| depth_rank | rank of sampling depth in the water column |  |
| sampling_date | sampling date |  |
| julian_day | julian day |  |
| longitude | longitude coordinates | degrees east |
| latitude | latitude coordinates | degrees north |
| OWD | days a given station was ice-free | days |
| by_OW_minus10_10 | classification of sectors based in OWD | sector |
| ice_concentration_percent | ice concentration cover | % |
| dna_concentration | dna concentration | ng.$\mu\text{L}^{-1}$ |
| dna_extraction_kit | dna extraction kit |  |
| n_reads | number of reads after filtering |  |
| reads_total | number of reads obtained from sequencing |  |
| pico_ml | pico-phytoplankton abundance | cells.mL$^{-1}$ |
| nano_ml | nano-phytoplankton abundance | cells.mL$^{-1}$ |
| pico_and_nano_ml | pico- and nano phytoplankton abundance | cells.mL$^{-1}$ |
| crypto_ml | cryptophyceae abundance | cells.mL$^{-1}$ |
| bact | bacteria abundance | cells.mL$^{-1}$ |
| temperature | temperature | degrees Celsius |
| fluo | fluorescence |  |
| cdom | colored dissolved organic matter | (ppb) |
| salinity | salinity |  |
| mixed_layer_depth | mixed layer depth | m |
| nitracline_depth | nitracline depth | m |
| PAR_irradiance | photosynthetically available radiation at 3 m depth | mol photons.m$^{-2}$.d$^{-1}$ |

**Table S1:** *(continued)*
| Variable | Decription | Unit |
| --- | --- | --- |
| primary_production | primary production | $\mu\text{gC.L}^{-1}.\text{day}^{-1}$ |
| primary_production_std_dev | primary production standard deviation | $\mu\text{gC.L}^{-1}.\text{day}^{-1}$ |
| don_release | dissolved organic nitrogen | $\text{nM.L}^{-1}.\text{day}^{-1}$ |
| NO3_assimilation | nitrate assimilation | $\text{nM.L}^{-1}.\text{day}^{-1}$ |
| NH4_assimilation | ammonium assimilation | $\text{nM.L}^{-1}.\text{day}^{-1}$ |
| urea_assimilation | urea assimilation | $\text{nM.L}^{-1}.\text{day}^{-1}$ |
| NH4_regeneration | ammonium regeneration | $\text{nM.L}^{-1}.\text{day}^{-1}$ |
| nitrification | nitrification | $\text{nM.L}^{-1}.\text{day}^{-1}$ |
| poc | particulate organic carbon | $\mu\text{M}$ |
| poc_std_dev | particulate organic carbon standard deviation | $\mu\text{M}$ |
| pon | particulate organic nitrogen | $\mu\text{M}$ |
| pon_std_dev | particulate organic nitrogen standard deviation | $\mu\text{M}$ |
| doc | dissolved organic carbon | $\mu\text{M}$ |
| don | dissolved organic nitrogen | $\mu\text{M}$ |
| nitrate | nitrate concentration | $\mu\text{M}$ |
| nitrite | nitrite concentration | $\mu\text{M}$ |
| phosphate | phosphate concentration | $\mu\text{M}$ |
| silica | orthosilicic acid concentration | $\mu\text{M}$ |
| ammonium | ammonium concentration | $\mu\text{M}$ |
| urea | urea concentration | $\mu\text{M}$ |
| ratio_NO3_SiOH4 | ratio nitrate to silica |  |
| ratio_PO4_NO3 | ratio phosphate to nitrate |  |
| ratio_NO3_PO4 | ratio nitrate to phosphate |  |
| chlorophyll_a | chlorophyll a concentration | $\text{mg.m}^{-3}$ |
| chlorophyll_b | chlorophyll b concentration | $\text{mg.m}^{-3}$ |

---

**Figure S1:**
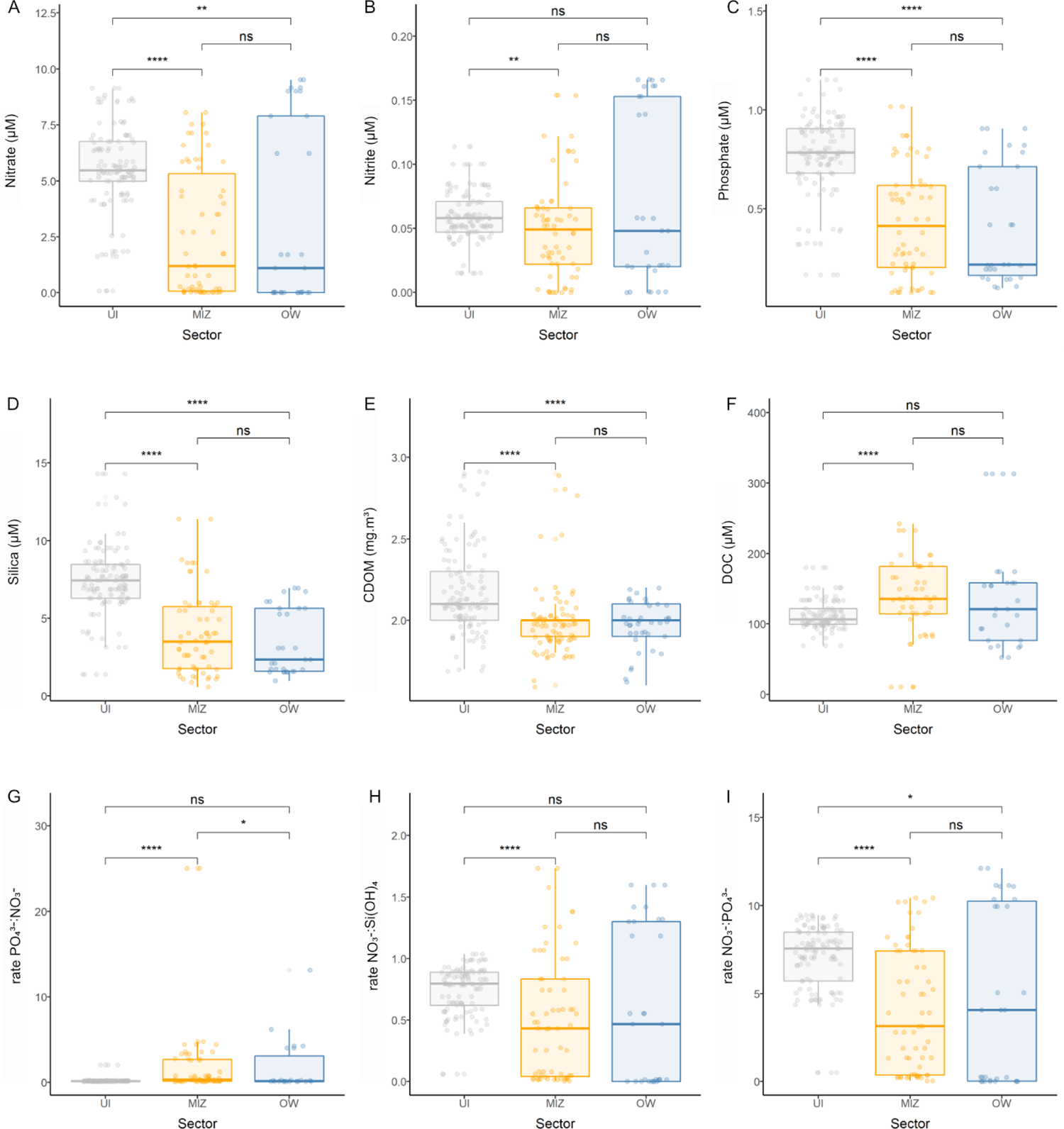
Nutrients for the three sectors: UI (grey), MIZ (yellow) and OW (blue); (A) nitrates (*µ*M); (B) nitrites (*µ*M); (C) phosphates (*µ*M); (D) orthosilicic acid (*µ*M); (E) colored dissolved organic matter (mg.m^3^); (F) dissolved organic carbon (*µ*M); (G) phosphate to nitrate ratio; (H) nitrate to orthosilicic acid ratio; (I) nitrate to phosphate ratio. Number of asterisks represent *p*-value obtained with the Wilcox test as follows: (*) *p ≤* 0.05; (**) *p ≤* 0.01; (***) *p ≤* 0.001; (****) *p ≤* 0.0001; “ns” = not significant.

**Figure S2:**
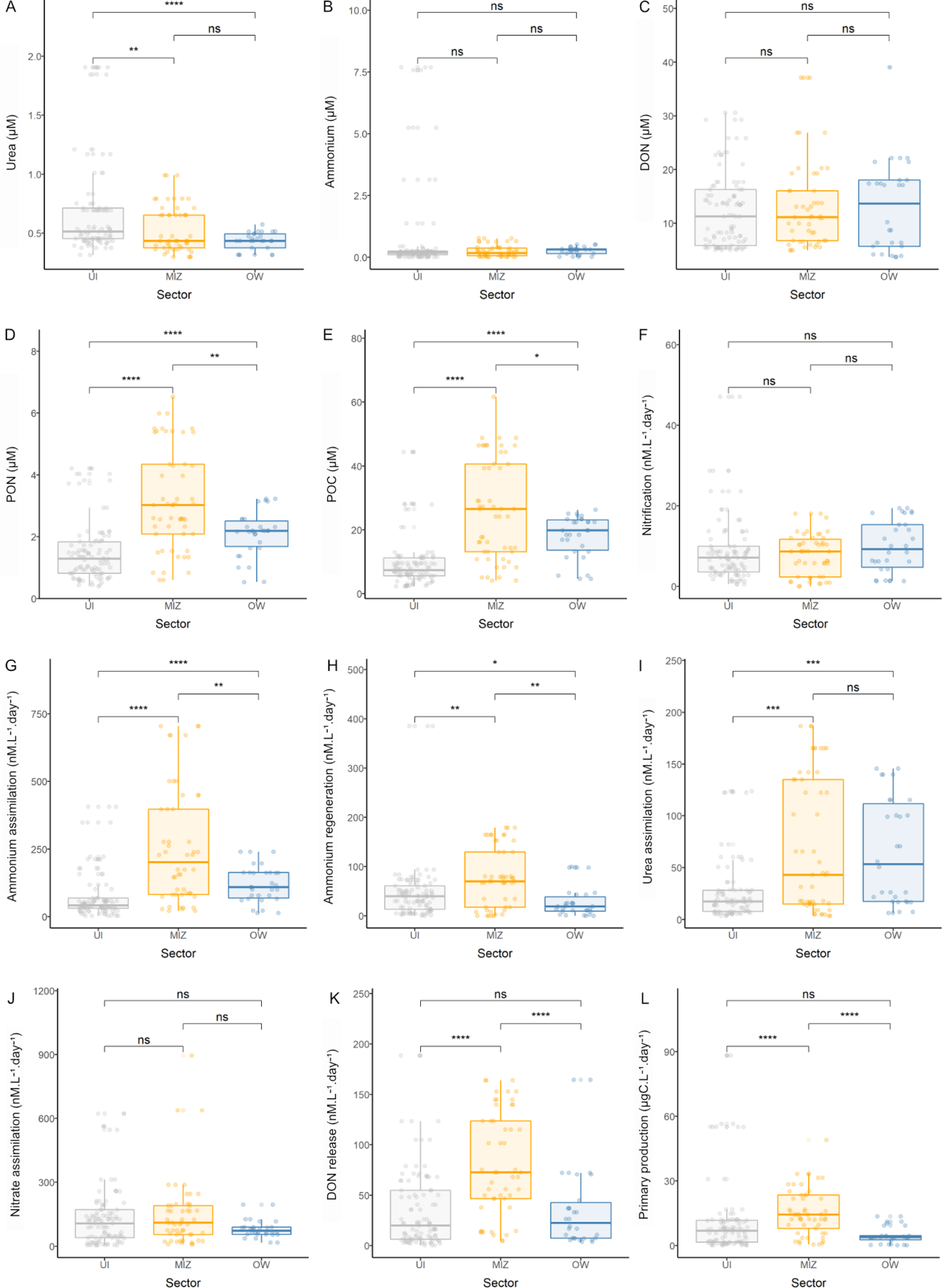
Nutrients and metabolic rates for the three sectors: UI (grey), MIZ (yellow) and OW (blue); (A) urea (*µ*M); (A) ammonium (*µ*M); (C) dissolved organic nitrogen (*µ*M); (D) particulate organic nitrogen (*µ*M); (E) particulate organic carbon (mg.m^3^); (F) nitrification (*µ*M); (G) ammonium assimilation (nM.L*^−^*^1^.day*^−^*^1^); (H) ammonium regeneration (nM.L*^−^*^1^.day*^−^*^1^); (I) urea assimilation (nM.L*^−^*^1^.day*^−^*^1^); (J) nitrate assimilation; (K) dissolved organic nitrogen (*µ*M); (L) primary production (*µ*gC.L*^−^*^1^.day*^−^*^1^). Number of asterisks represent *p*-value obtained with the Wilcox test as follows: (*) *p ≤* 0.05; (**) *p ≤* 0.01; (***) *p ≤* 0.001; (****) *p ≤* 0.0001; “ns” = not significant.

**Figure S3:**
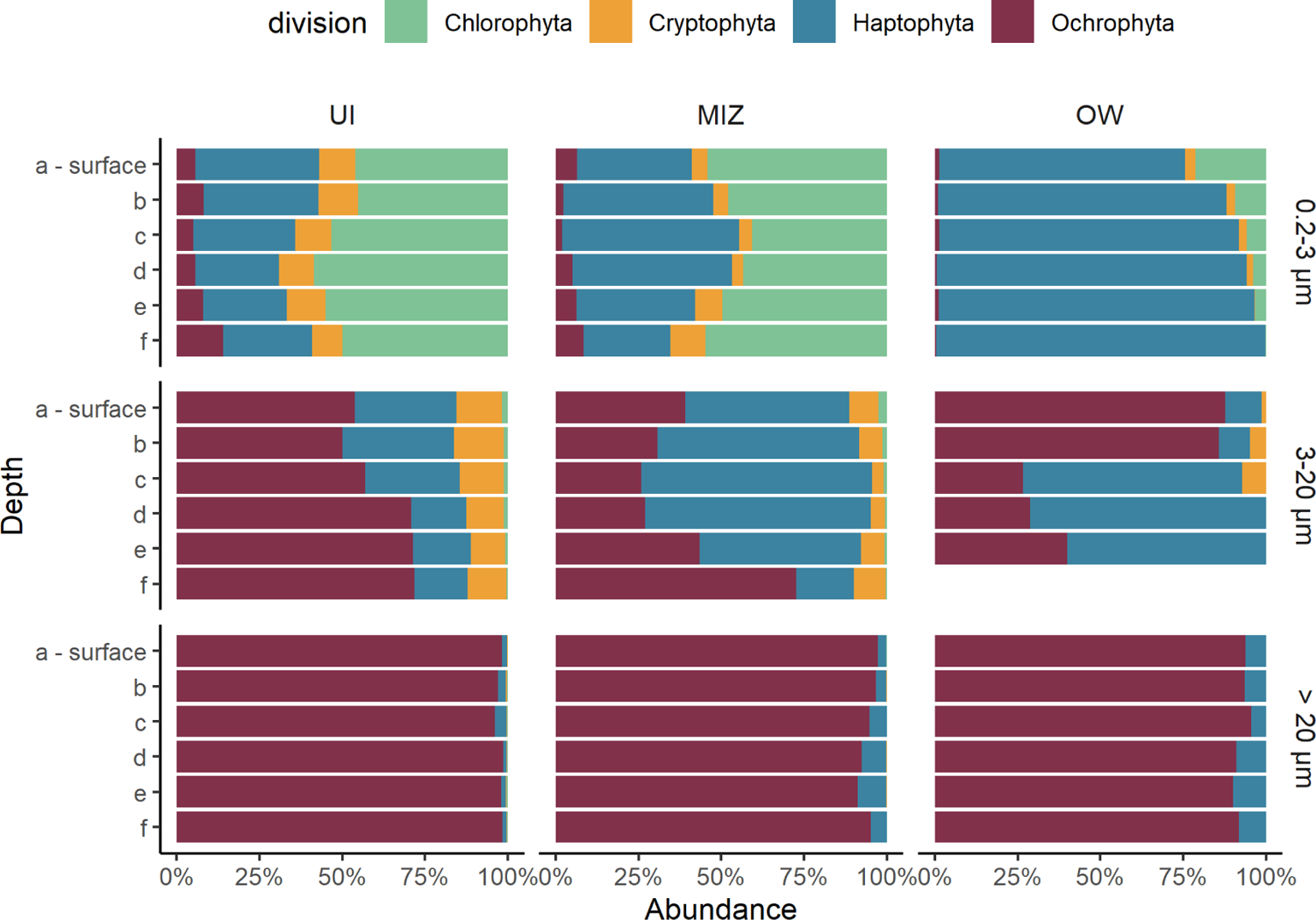
Relative abundance of reads at the division level between sectors and size fractions. UI: Under Ice; MIZ: Marginal Ice Zone; OW: Open Water; letters on the y-axis refer to the depth level where “a” corresponds to the surface and “f” to the deepest sampled depth, usually between 40 m and 60 m.

**Figure S4:**
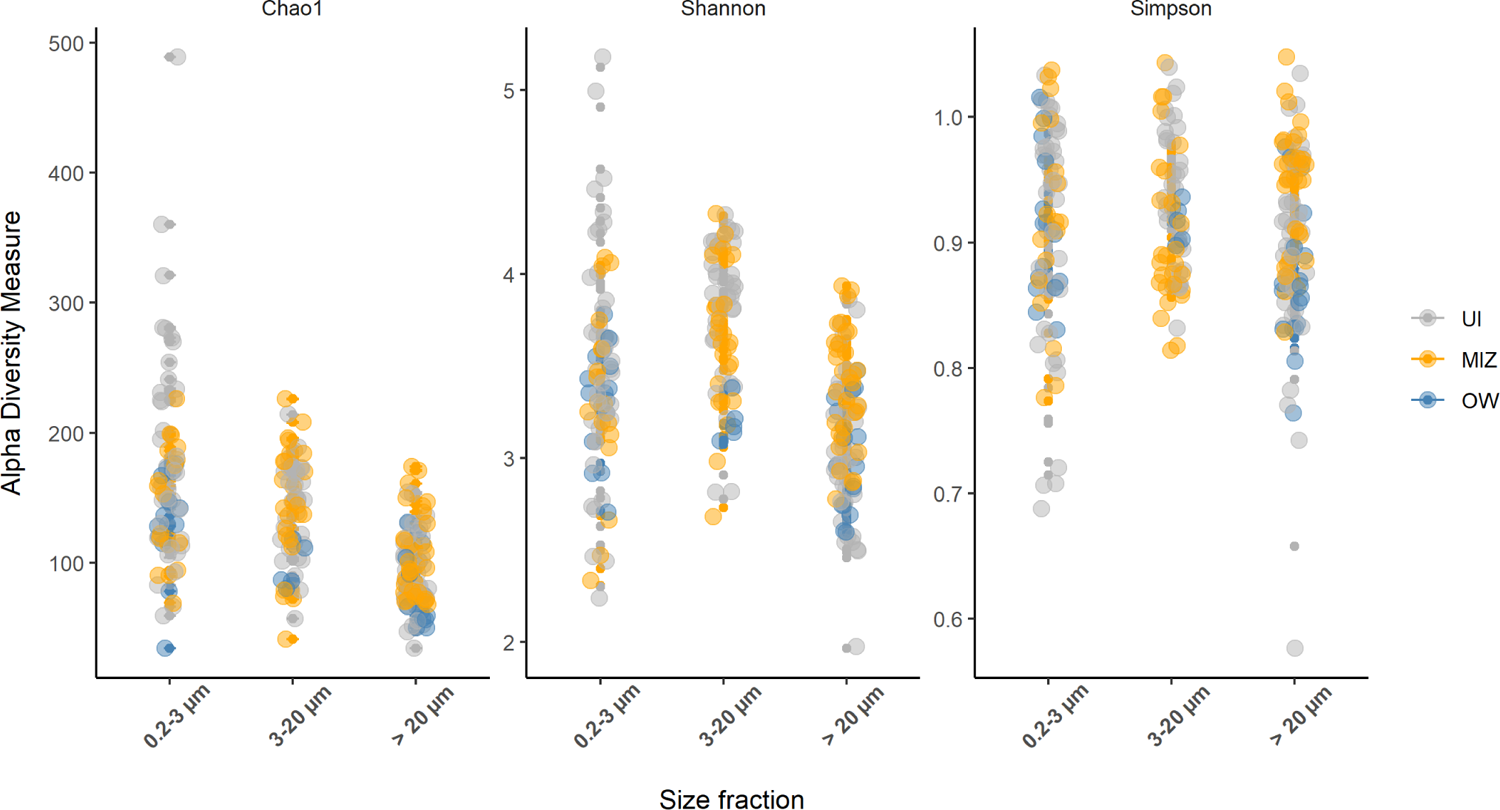
Chao1, Shannon and Simpson alpha diversity indices divided by size fraction; sectors are represented by the colors grey (UI), yellow (MIZ) and blue (OW).

**Figure S5:**
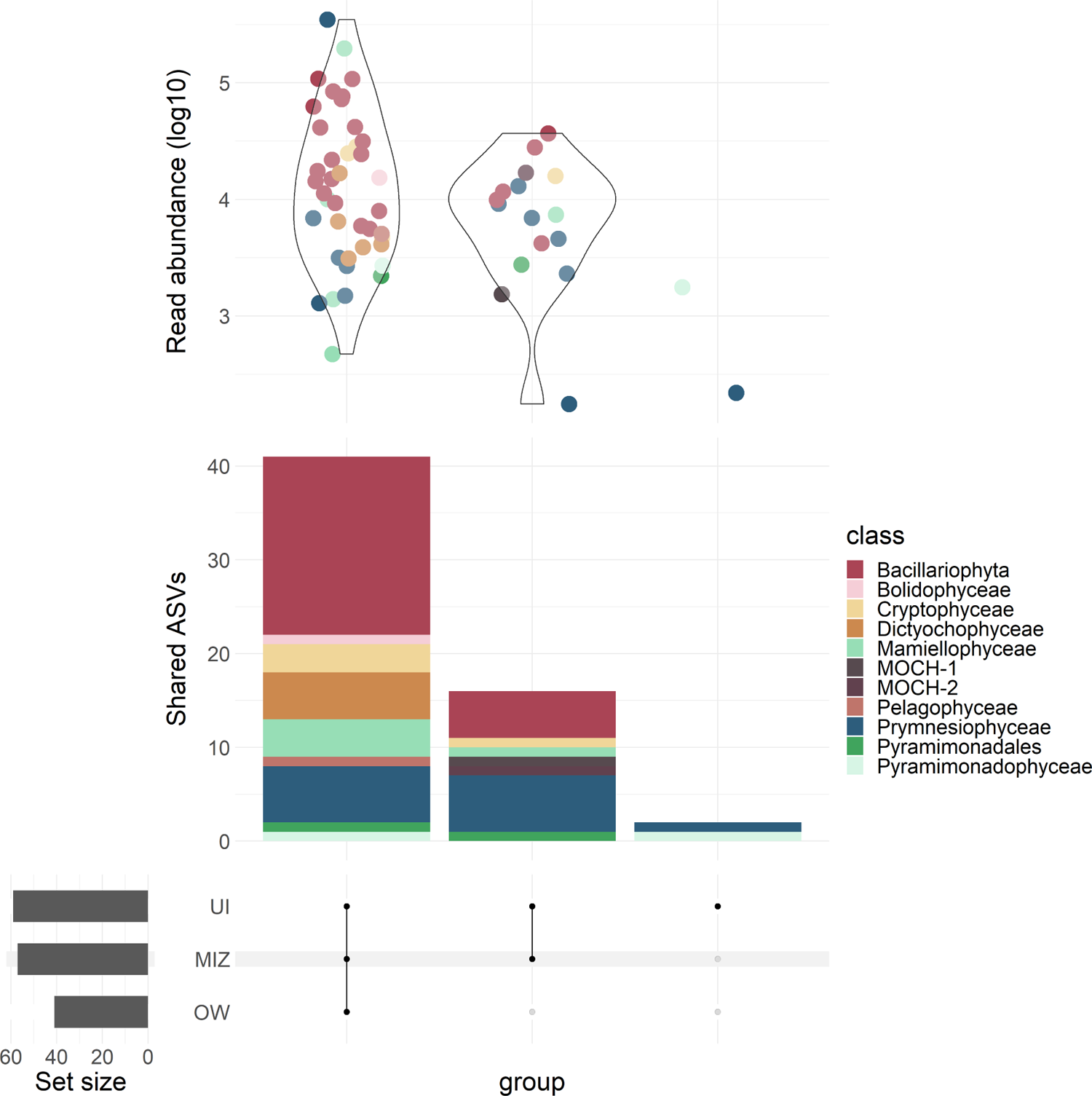
Number of ASVs from the abundant community exclusive from or shared between the sectors UI (under ice), MIZ (marginal ice-zone), and OW (open water); colors represent the class from each ASV; read abundance (in log10) is displayed at the top of each intersection; the names and assignations of the ASVs exclusive from ice-associated sectors are shown in grey panels.

**Figure S6:**
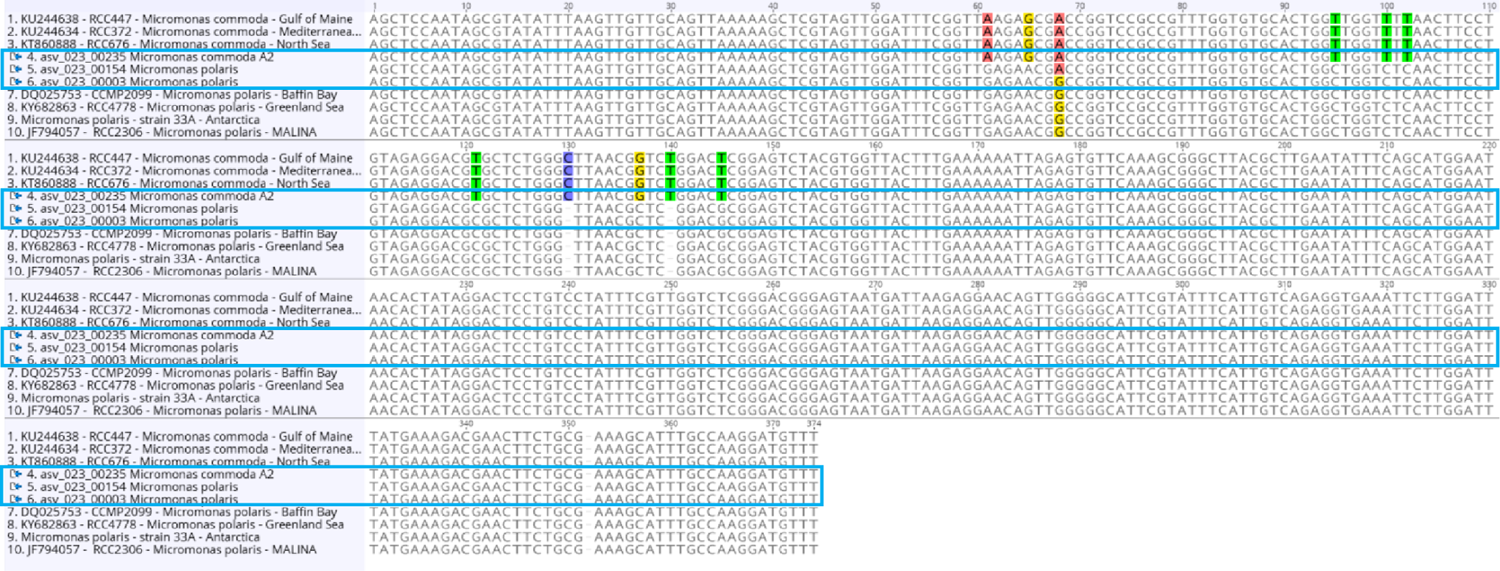
Sequence alignment of the 18S rRNA of *Micromonas* ASVs showing two *M. polaris* ASVs with a single nucleotide difference, and a *M. commoda* A2 ASV.

**Figure S7:**
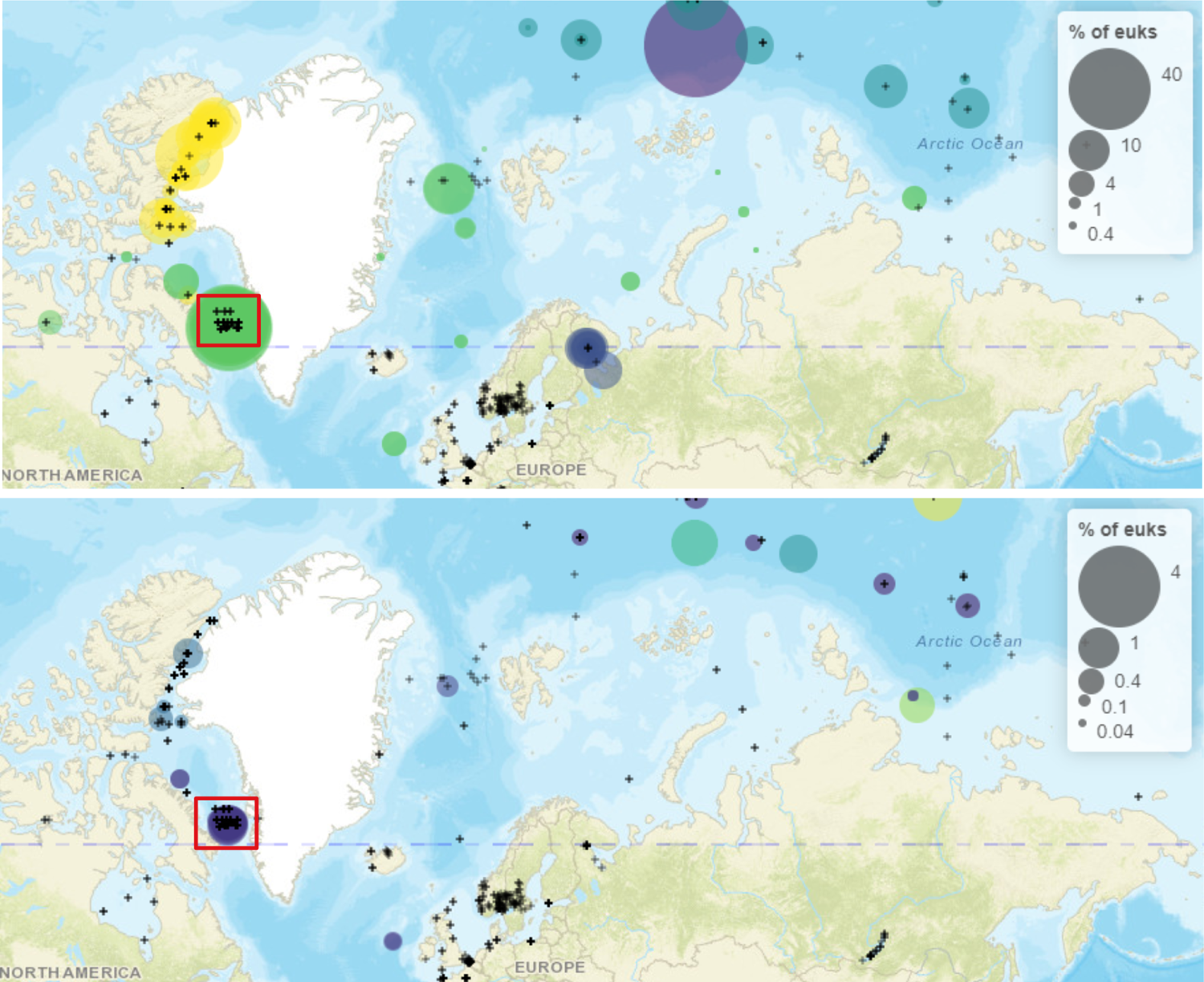
Partial snapshot of *M. polaris* ASV_0003 (top panel) and ASV_0154 (lower panel) distribution in the metaPR^2^ database showing 100% similar reads from other studies. Colors indicate different sampling campaigns within metaPR^2^. Size of bubbles represent the percentage in relation to other eukaryotes within each station. Note that maximum percentages are distinct between panels to compensate for the lower abundance of ASV_0154.

**Figure S8:**
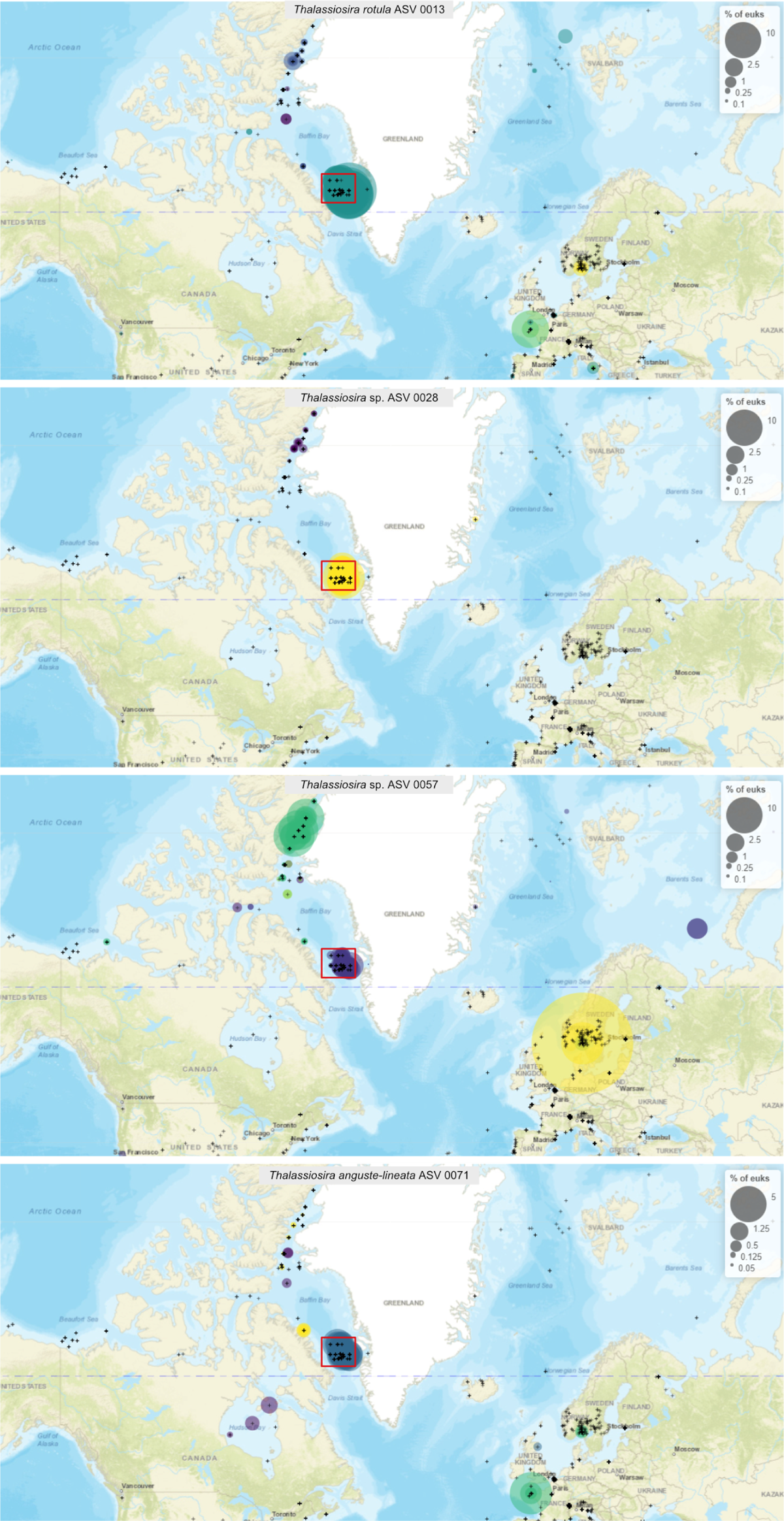
Partial snapshot of *Thalassiosira* ASV_0013 (top panel), ASV_0057 (middle panel), and ASV_0071 (lower panel) distribution in the metaPR^2^ database showing 100% similar reads from other studies. Colors indicate different sampling campaigns within metaPR^2^. Size of bubbles represent the percentage in relation to other eukaryotes within each station. Note that maximum percentages are distinct between panels. The approximate region of sampling from the present study is marked by a red square.

## Notes

https://github.com/catherine-gerikas/GE_Amundsen_18S_metaB_supplementary_material

### Competing Interest Statement

The authors have declared no competing interest.

## References cited

1. Alou-Font, E., Roy, S., Agustí, S., & Gosselin, M. (2016). Cell viability, pigments and photosynthetic performance of Arctic phytoplankton in contrasting ice-covered and open-water conditions during the spring-summer transition. Marine Ecology Progress Series, 543, 89–106. https://doi.org/10.3354/meps11562

2. Ardyna, M., Mundy, C. J., Mayot, N., Matthes, L. C., Oziel, L., Horvat, C., Leu, E., Assmy, P., Hill, V., Matrai, P. A., Gale, M., Melnikov, I. A., & Arrigo, K. R. (2020). Under-Ice Phytoplankton Blooms: Shedding Light on the “Invisible” Part of Arctic Primary Production. Frontiers in Marine Science, 7, 1–25. https://doi.org/10.3389/fmars.2020.608032

3. Arrigo, K. R., Perovich, D. K., Pickart, R. S., Brown, Z. W., van Dijken, G. L., Lowry, K. E., Mills, M. M., Palmer, M. A., Balch, W. M., Bahr, F., Bates, N. R., Benitez-Nelson, C., Bowler, B., Brownlee, E., Ehn, J. K., Frey, K. E., Garley, R., Laney, S. R., Lubelczyk, L., … Swift, J. H. (2012). Massive Phytoplankton Blooms Under Arctic Sea Ice. Science, 336, 1408–1408. https://doi.org/10.1126/science.1215065

4. Arrigo, K. R., Perovich, D. K., Pickart, R. S., Brown, Z. W., van Dijken, G. L., Lowry, K. E., Mills, M. M., Palmer, M. A., Balch, W. M., Bates, N. R., Benitez-Nelson, C. R., Brownlee, E., Frey, K. E., Laney, S. R., Mathis, J., Matsuoka, A., Greg Mitchell, B., Moore, G. W., Reynolds, R. A., … Swift, J. H. (2014). Phytoplankton blooms beneath the sea ice in the Chukchi sea. Deep-Sea Research Part II: Topical Studies in Oceanography, 105, 1–16. https://doi.org/10.1016/j.dsr2.2014.03.018

5. Assmy, P., Fernández-Méndez, M., Duarte, P., Meyer, A., Randelhoff, A., Mundy, C. J., Olsen, L. M., Kauko, H. M., Bailey, A., Chierici, M., Cohen, L., Doulgeris, A. P., Ehn, J. K., Fransson, A., Gerland, S., Hop, H., Hudson, S. R., Hughes, N., Itkin, P., … Granskog, M. A. (2017). Leads in Arctic pack ice enable early phytoplankton blooms below snow-covered sea ice. Scientific Reports, 7, 1–9. https://doi.org/10.1038/srep40850

6. Balzano, S., Marie, D., Gourvil, P., & Vaulot, D. (2012). Composition of the summer photosynthetic pico and nanoplankton communities in the Beaufort Sea assessed by T-RFLP and sequences of the 18S rRNA gene from flow cytometry sorted samples. The ISME journal, 6, 1480–1498. https://doi.org/10.1038/ismej.2011.213

7. Balzano, S., Percopo, I., Siano, R., Gourvil, P., Chanoine, M., Marie, D., Vaulot, D., & Sarno, D. (2017). Morphological and genetic diversity of Beaufort Sea diatoms with high contributions from the *Chaetoceros neogracilis* species complex. Journal of Phycology, 53, 161–187. https://doi.org/10.1111/jpy.12489

8. Benner, I., Irwin, A. J., & Finkel, Z. (2019). Capacity of the common Arctic picoeukaryote *Micromonas* to adapt to a warming warming ocean. Limnology and Oceanography Letters, 5, 221–227. https://doi.org/10.6084/m9

9. Bluhm, B. A., Kosobokova, K. N., & Carmack, E. C. (2015). A tale of two basins: An integrated physical and biological perspective of the deep Arctic Ocean. Progress in Oceanography, 139, 89–121. https://doi.org/10.1016/j.pocean.2015.07.011

10. Boetius, A., Albrecht, S., Bakker, K., Bienhold, C., Felden, J., Fernández-Méndez, M., Hendricks, S., Katlein, C., Lalande, C., Krumpen, T., Nicolaus, M., Peeken, I., Rabe, B., Rogacheva, A., Rybakova, E., Somavilla, R., Wenzhöfer, F., & Felden, J. (2013). Export of algal biomass from the melting arctic sea ice. Science, 339, 1430–1432. https://doi.org/10.1126/science.1231346

11. Bruyant, F., Amiraux, R., Amyot, M.-P., Archambault, P., Artigue, L., Bardedo de Freitas, L., Bécu, G., Bélanger, S., Bourgain, P., Bricaud, A., Brouard, E., Brunet, C., Burgers, T., Caleb, D., Chalut, K., Clautre, H., Cornet-Barthaux, V., Coupel, P., Cusa, M., … Babin, M. (2022). The Green Edge cruise: Understanding the onset, life and fate of the Arctic phytoplankton spring bloom. Earth Syst. Sci., preprint, 1–47.

12. Callahan, B. J., McMurdie, P. J., Rosen, M. J., Han, A. W., Johnson, A. J. A., & Holmes, S. P. (2016). DADA2: High-resolution sample inference from Illumina amplicon data. Nature Methods, 13, 581–583. https://doi.org/10.1038/nmeth.3869

13. Chang, F. H., Sutherland, J., Ehrenberg, D., & Schrader, D. (2017). Taxonomic revision of Dictyochales (Dicty-ochophyceae) based on morphological, ultrastructural, biochemical and molecular data. Phycological Research, 65, 235–247. https://doi.org/10.1111/pre.12181

14. Comeau, A. M., Philippe, B., Thaler, M., Gosselin, M., Poulin, M., & Lovejoy, C. (2013). Protists in Arctic Drift and Land-Fast Sea Ice. Journal of Phycology, 49, 229–240. https://doi.org/10.1111/jpy.12026

15. Crawford, D. W., Cefarelli, A. O., Wrohan, I. A., Wyatt, S. N., & Varela, D. E. (2018). Spatial patterns in abundance, taxonomic composition and carbon biomass of nano- and microphytoplankton in Subarctic and Arctic Seas. Progress in Oceanography, 162, 132–159. https://doi.org/10.1016/j.pocean.2018.01.006

16. De Cáceres, M., Legendre, P., & Moretti, M. (2010). Improving indicator species analysis by combining groups of sites. Oikos, 119, 1674–1684. https://doi.org/10.1111/j.1600-0706.2010.18334.x

17. Demory, D., Baudoux, A.-C., Monier, A., Simon, N., Six, C., Ge, P., Marie, D., Sciandra, A., & Rabouille, S. (2019). Picoeukaryotes of the *Micromonas* genus: sentinels of a warming ocean. The ISME Journal, 13, 132–146.

18. Dixon, P. (2003). Computer program review VEGAN, a package of R functions for community ecology. Journal of Vegetation Science, 14, 927–930. http://doi.wiley.com/10.1111/j.1654-1103.2002.tb02049.x

19. Egge, E. S., Eikrem, W., & Edvardsen, B. (2014). Deep-branching Novel Lineages and High Diversity of Haptophytes in the Skagerrak (Norway) Uncovered by 454 pyrosequencing. The Journal of Eukaryotic Microbiology, 1–20. https://doi.org/10.1111/jeu.12157

20. Freyria, N. J., Joli, N., & Lovejoy, C. (2021). A decadal perspective on north water microbial eukaryotes as Arctic Ocean sentinels. Scientific Reports, 11, 1–14. https://doi.org/10.1038/s41598-021-87906-4

21. García-García, N., Tamames, J., Linz, A. M., Pedrós-Alió, C., & Puente-Sánchez, F. (2019). Microdiversity ensures the maintenance of functional microbial communities under changing environmental conditions. ISME Journal, 13, 2969–2983. https://doi.org/10.1038/s41396-019-0487-8

22. Gradinger, R. (1996). Occurrence of an algal bloom under Arctic pack ice. Marine Ecology Progress Series, 131, 301–305. https://doi.org/10.3354/meps131301

23. Guillou, L., Bachar, D., Audic, S., Bass, D., Berney, C., Bittner, L., Boutte, C., Burgaud, G., De Vargas, C., Decelle, J., Del Campo, J., Dolan, J. R., Dunthorn, M., Edvardsen, B., Holzmann, M., Kooistra, W. H., Lara, E., Le Bescot, N., Logares, R., … Christen, R. (2013). The Protist Ribosomal Reference database (PR^2^): A catalog of unicellular eukaryote Small Sub-Unit rRNA sequences with curated taxonomy. Nucleic Acids Research, 41, D597–D604. https://doi.org/10.1093/nar/gks1160

24. Hardge, K., Peeken, I., Neuhaus, S., Lange, B. A., Stock, A., Stoeck, T., Weinisch, L., & Metfies, K. (2017). The importance of sea ice for exchange of habitat-specific protist communities in the Central Arctic Ocean. Journal of Marine Systems, 165, 124–138. https://doi.org/10.1016/j.jmarsys.2016.10.004

25. Hegseth, N. E., & Sundfjord, A. (2008). Intrusion and blooming of Atlantic phytoplankton species in the high Arctic. Journal of Marine Systems, 74, 108–119. https://doi.org/10.1016/j.jmarsys.2007.11.011

26. Hilligsøe, K. M., Richardson, K., Bendtsen, J., Sørensen, L. L., Nielsen, T. G., & Lyngsgaard, M. M. (2011). Linking phytoplankton community size composition with temperature, plankton food web structure and sea-air CO2 flux. Deep-Sea Research Part I: Oceanographic Research Papers, 58, 826–838. https://doi.org/10.1016/j.dsr.2011.06.004

27. Hop, H., Mundy, C. J., Gosselin, M., Rossnagel, A. L., & Barber, D. G. (2011). Zooplankton boom and ice amphipod bust below melting sea ice in the Amundsen Gulf, Arctic Canada. Polar Biology, 34, 1947– 1958. https://doi.org/10.1007/s00300-011-0991-4

28. Hop, H., Vihtakari, M., Bluhm, B. A., Assmy, P., Poulin, M., Gradinger, R., Peeken, I., von Quillfeldt, C., Olsen, L. M., Zhitina, L., & Melnikov, I. A. (2020). Changes in Sea-Ice Protist Diversity With Declining Sea Ice in the Arctic Ocean From the 1980s to 2010s. Frontiers in Marine Science, 7. https://doi.org/10.3389/fmars.2020.00243

29. Hoppe, C. J. M., Flintrop, C. M., & Rost, B. (2018). The arctic picoeukaryote *Micromonas pusilla* benefits synergistically from warming and ocean acidification. Biogeosciences, 15, 4353–4365. https://doi.org/10.5194/bg-15-4353-2018

30. Horner, R., Ackley, S. F., Dieckmann, G. S., Gulliksen, B., Hoshiai, T., Legendre, L., Melnikov, I. A., Reeburgh, W. S., Spindler, M., & Sullivan, C. W. (1992). Ecology of sea ice biota − 1. Habitat, terminology, and methodology. Polar Biology, 12, 417–427. https://doi.org/10.1007/BF00243113

31. Horvat, C., Jones, D. R., Iams, S., Schroeder, D., Flocco, D., & Feltham, D. (2017). The frequency and extent of sub-ice phytoplankton blooms in the Arctic Ocean. Science Advances, 3, e1601191. https://doi.org/10.1126/sciadv.1601191

32. Janout, M. A., Hölemann, J., Waite, A. M., Krumpen, T., von Appen, W. J., & Martynov, F. (2016). Sea-ice retreat controls timing of summer plankton blooms in the Eastern Arctic Ocean. Geophysical Research Letters, 43, 12, 493–12, 501. https://doi.org/10.1002/2016GL071232

33. Joli, N., Monier, A., Logares, R., & Lovejoy, C. (2017). Seasonal patterns in Arctic prasinophytes and inferred ecology of Bathycoccus unveiled in an Arctic winter metagenome. ISME Journal, 11, 1372–1385. https://doi.org/10.1038/ismej.2017.7

34. Jones, E. P., Swift, J. H., Anderson, L. G., Lipizer, M., Civitarese, G., Falkner, K. K., Kattner, G., & McLaughlin, F. (2003). Tracing Pacific water in the North Atlantic Ocean. Journal of Geophysical Research C: Oceans, 108, 13–1. https://doi.org/10.1029/2001jc001141

35. Kalenitchenko, D., Joli, N., Potvin, M., Tremblay, J. é., & Lovejoy, C. (2019). Biodiversity and species change in the Arctic Ocean: A view through the lens of Nares Strait. Frontiers in Marine Science, 6, 1–17. https://doi.org/10.3389/fmars.2019.00479

36. Kauko, H. M., Olsen, L. M., Duarte, P., Peeken, I., Granskog, M. A., Johnsen, G., Fernández-Méndez, M., Pavlov, A. K., Mundy, C. J., & Assmy, P. (2018). Algal Colonization of Young Arctic Sea Ice in Spring. Frontiers in Marine Science, 5, 1–20. https://doi.org/10.3389/fmars.2018.00199

37. Kinney, J. C., Maslowski, W., Osinski, R., Jin, M., Frants, M., Jeffery, N., & Lee, Y. J. (2020). Hidden Production: On the Importance of Pelagic Phytoplankton Blooms Beneath Arctic Sea Ice. Journal of Geophysical Research: Oceans, 125. https://doi.org/10.1029/2020JC016211

38. Kopf, A., Bicak, M., Kottmann, R., Schnetzer, J., Kostadinov, I., Lehmann, K., Fernandez-Guerra, A., Jeanthon, C., Rahav, E., Ullrich, M., Wichels, A., Gerdts, G., Polymenakou, P., Kotoulas, G., Siam, R., Abdallah, R. Z., Sonnenschein, E. C., Cariou, T., O’Gara, F., … Glöckner, F. O. (2015). The ocean sampling day consortium. GigaScience, 4. https://doi.org/10.1186/s13742-015-0066-5

39. Krassowski, M. (2020). Complexupset. https://doi.org/10.5281/zenodo.3700590

40. Kvernvik, A. C., Rokitta, S. D., Leu, E., Harms, L., Gabrielsen, T. M., Rost, B., & Hoppe, C. J. (2020). Higher sensitivity towards light stress and ocean acidification in an Arctic sea-ice-associated diatom compared to a pelagic diatom. New Phytologist, 226, 1708–1724. https://doi.org/10.1111/nph.16501

41. Lafond, A., Leblanc, K., Quéguiner, B., Moriceau, B., Leynaert, A., Cornet, V., Legras, J., Ras, J., Parenteau, M., Garcia, N., Babin, M., & Tremblay, J.-É. (2019). Late spring bloom development of pelagic diatoms in Baffin Bay. Elementa: Science of the Anthropocene, 7, 1–24.

42. Lalande, C., Bauerfeind, E., Nöthig, E. M., & Beszczynska-Möller, A. (2013). Impact of a warm anomaly on export fluxes of biogenic matter in the eastern Fram Strait. Progress in Oceanography, 109, 70–77. https://doi.org/10.1016/j.pocean.2012.09.006

43. Lasternas, S., & Agustı, S. (2010). Phytoplankton community structure during the record Arctic ice-melting of summer 2007. Polar Biology, 33, 1709–1717. https://doi.org/10.1007/s00300-010-0877-x

44. Leu, E., Mundy, C. J., Assmy, P., Campbell, K., Gabrielsen, T. M., Gosselin, M., Juul-Pedersen, T., & Gradinger, R. (2015). Arctic spring awakening - Steering principles behind the phenology of vernal ice algal blooms. Progress in Oceanography, 139, 151–170. https://doi.org/10.1016/j.pocean.2015.07.012

45. Leu, E., Søreide, J. E., Hessen, D. O., Falk-Petersen, S., & Berge, J. (2011). Consequences of changing sea-ice cover for primary and secondary producers in the European Arctic shelf seas: Timing, quantity, and quality. Progress in Oceanography, 90, 18–32. https://doi.org/10.1016/j.pocean.2011.02.004

46. Lewis, K. M., Arntsen, A. E., Coupel, P., Joy-Warren, H., Lowry, K. E., Matsuoka, A., Mills, M. M., van Dijken, G. L., Selz, V., & Arrigo, K. R. (2019). Photoacclimation of Arctic Ocean phytoplankton to shifting light and nutrient limitation. Limnology and Oceanography, 64, 284–301. https://doi.org/10.1002/lno.11039

47. Li, W. K., McLaughlin, F. A., Lovejoy, C., & Carmack, E. C. (2009). Smallest algae thrive as the Arctic Ocean freshens. Science, 326, 539. https://doi.org/10.1126/science.1179798

48. Lovejoy, C., Legendre, L., Martineau, M. J., Bâcle, J., & Von Quillfeldt, C. H. (2002). Distribution of phytoplankton and other protists in the North Water. Deep-Sea Research Part II: Topical Studies in Oceanography, 49, 5027–5047. https://doi.org/10.1016/S0967-0645(02)00176-5

49. Lovejoy, C., & Potvin, M. (2011). Microbial eukaryotic distribution in a dynamic Beaufort Sea and the Arctic Ocean. Journal of Plankton Research, 33, 431–444. https://doi.org/10.1093/plankt/fbq124

50. Lovejoy, C., Vincent, W. F., Bonilla, S., Roy, S., Martineau, M. J., Terrado, R., Potvin, M., Massana, R., & Pedrós-Alió, C. (2007). Distribution, phylogeny, and growth of cold-adapted picoprasinophytes in arctic seas. Journal of Phycology, 43, 78–89. https://doi.org/10.1111/j.1529-8817.2006.00310.x

51. Luo, W., Li, H., Gao, S., Yu, Y., Lin, L., & Zeng, Y. (2016). Molecular diversity of microbial eukaryotes in sea water from Fildes Peninsula, King George Island, Antarctica. Polar Biology, 39, 605–616. https://doi.org/10.1007/s00300-015-1815-8

52. Marie, D., Partensky, F., Jacquet, S., & Vaulot, D. (1997). Enumeration and cell cycle analysis of natural populations of marine picoplankton by flow cytometry using the nucleic acid stain SYBR Green I. Applied and Environmental Microbiology, 63, 186–193. https://doi.org/10.1128/aem.63.1.186-193.1997

53. Marie, D., Shi, X. L., Rigaut-Jalabert, F., & Vaulot, D. (2010). Use of flow cytometric sorting to better assess the diversity of small photosynthetic eukaryotes in the English Channel. FEMS microbiology ecology, 72, 165–78. https://doi.org/10.1111/j.1574-6941.2010.00842.x

54. Martin, J., Tremblay, J. É., Gagnon, J., Tremblay, G., Lapoussière, A., Jose, C., Poulin, M., Gosselin, M., Gratton, Y., & Michel, C. (2010). Prevalence, structure and properties of subsurface chlorophyll maxima in Canadian Arctic waters. Marine Ecology Progress Series, 412, 69–84. https://doi.org/10.3354/meps08666

55. Massicotte, P., Amiraux, R., Amyot, M. P., Archambault, P., Ardyna, M., Arnaud, L., Artigue, L., Aubry, C., Ayotte, P., Bécu, G., Bélanger, S., Benner, R., Bittig, H. C., Bricaud, A., Brossier, É., Bruyant, F., Chauvaud, L., Christiansen-Stowe, D., Claustre, H., … Babin, M. (2020). Green edge ice camp campaigns: Understanding the processes controlling the under-ice arctic phytoplankton spring bloom. Earth System Science Data, 12, 151–176. https://doi.org/10.5194/essd-12-151-2020

56. Matsuoka, A., Larouche, P., Poulin, M., Vincent, W., & Hattori, H. (2009). Phytoplankton community adaptation to changing light levels in the southern Beaufort Sea, Canadian Arctic. Estuarine, Coastal and Shelf Science, 82, 537–546. https://doi.org/10.1016/j.ecss.2009.02.024

57. McMurdie, P. J., & Holmes, S. (2013). Phyloseq: An R Package for Reproducible Interactive Analysis and Graphics of Microbiome Census Data. PLoS ONE, 8. https://doi.org/10.1371/journal.pone.0061217

58. Meredith, M., Sommerkorn, M., Cassotta, S., Derksen, C., Ekaykin, A., Hollowed, A., Kofinas, G., Mackintosh, A., Melbourne-Thomas, J., Muelbert, M., Ottersen, G., Pritchard, H., & Schuur, E. (2019). Polar regions, In Ipcc special report on the ocean and cryosphere in a changing climate. https://doi.org/10.1016/S1366-7017(01)00066-6

59. Metfies, K., Von Appen, W. J., Kilias, E., Nicolaus, A., & Nöthig, E. M. (2016). Biogeography and photosynthetic biomass of arctic marine pico-eukaroytes during summer of the record sea ice minimum 2012. PLoS ONE, 11, 1–20. https://doi.org/10.1371/journal.pone.0148512

60. Moestrup, Ø., & Thomsen, H. (1990). *DICTYOCHA SPECULUM* (SILICOFLAGELLATA, DICTYOCHOPHYCEAE). STUDIES ON ARMOURED AND UNARMOURED STAGES. Danske Vidensk. Selsk. Biol. Skr., 37, 1–56.

61. Morán, X. A. G., López-Urrutia, Á., Calvo-Díaz, A., & Li, W. K. (2010). Increasing importance of small phytoplankton in a warmer ocean. Global Change Biology, 16, 1137–1144. https://doi.org/10.1111/j.1365-2486.2009.01960.x

62. Morando, M., & Capone, D. G. (2018). Direct utilization of organic nitrogen by phytoplankton and its role in nitrogen cycling within the southern California bight. Frontiers in Microbiology, 9, 1–13. https://doi.org/10.3389/fmicb.2018.02118

63. Münchow, A., Falkner, K. K., & Melling, H. (2015). Baffin Island and West Greenland Current Systems in northern Baffin Bay. Progress in Oceanography, 132, 305–317. https://doi.org/10.1016/j.pocean.2014.04.001

64. Mundy, C. J., Gosselin, M., Ehn, J. K., Belzile, C., Poulin, M., Alou, E., Roy, S., Hop, H., Lessard, S., Papakyriakou, T. N., Barber, D. G., & Stewart, J. (2011). Characteristics of two distinct high-light acclimated algal communities during advanced stages of sea ice melt. Polar Biology, 34, 1869–1886. https://doi.org/10.1007/s00300-011-0998-x

65. Needham, D. M., & Fuhrman, J. A. (2016). Pronounced daily succession of phytoplankton, archaea and bacteria following a spring bloom. Nature microbiology, 1, 1–7. https://doi.org/10.1038/nmicrobiol.2016.5

66. Neukermans, G., Oziel, L., & Babin, M. (2018). Increased intrusion of warming Atlantic waters leads to rapid expansion of temperate phytoplankton in the Arctic. Global Change Biology, 1–9. https://doi.org/10.1111/gcb.14075

67. Niemi, A., Michel, C., Hille, K., & Poulin, M. (2011). Protist assemblages in winter sea ice: setting the stage for the spring ice algal bloom. Polar Biology, 34, 1803–1817. https://doi.org/10.1007/s00300-011-1059-1

68. Not, F., Massana, R., Latasa, M., Marie, D., Pedro, C., Vaulot, D., & Simon, N. (2005). Late summer community composition and abundance of photosynthetic picoeukaryotes in Norwegian and Barents Seas. Limnology and Oceanography, 50, 1677–1686.

69. Olsen, L. M., Laney, S. R., Duarte, P., Kauko, H. M., Fernández-méndez, M., Mundy, C. J., Rösel, A., Meyer, A., Itkin, P., Cohen, L., Peeken, I., Tatarek, A., Róźańska-Pluta, M., Wiktor, J., Taskjelle, T., Pavlov, A. K., Hudson, S. R., Granskog, M. A., Hop, H., & Assmy, P. (2017). The seeding of ice algal blooms in Arctic pack ice: The multiyear ice seed repository hypothesis. Journal of Geophysical Research: Biogeosciences, 122, 1529–1548. https://doi.org/10.1002/2016JG003668

70. Oziel, L., Baudena, A., Ardyna, M., Massicotte, P., Randelhoff, A., Sallée, J. B., Ingvaldsen, R. B., Devred, E., & Babin, M. (2020). Faster Atlantic currents drive poleward expansion of temperate phytoplankton in the Arctic Ocean. Nature Communications, 11. https://doi.org/10.1038/s41467-020-15485-5

71. Oziel, L., Massicotte, P., Randelhoff, A., Ferland, J., Vladoiu, A., Lacour, L., Galindo, V., Lambert-Girard, S., Dumont, D., Cuypers, Y., Bouruet-Aubertot, P., Mundy, C.-J., Ehn, J., Bécu, G., Marec, C., Forget, M.-H., Garcia, N., Coupel, P., Raimbault, P., … Babin, M. (2019). Environmental factors influencing the seasonal dynamics of spring algal blooms in and beneath sea ice in western Baffin Bay. Elementa: Science of the Anthropocene, 7, 34. https://doi.org/10.1525/elementa.372

72. Perrette, M., Yool, A., Quartly, G. D., & Popova, E. E. (2011). Near-ubiquity of ice-edge blooms in the Arctic. Biogeosciences, 8, 515–524. https://doi.org/10.5194/bg-8-515-2011

73. Piredda, R., Tomasino, M. P., D’Erchia, A. M., Manzari, C., Pesole, G., Montresor, M., Kooistra, W. H., Sarno, D., & Zingone, A. (2017). Diversity and temporal patterns of planktonic protist assemblages at a Mediterranean Long Term Ecological Research site. FEMS Microbiology Ecology, 93, 1–14. https://doi.org/10.1093/femsec/fiw200

74. Polyakov, I. V., Pnyushkov, A. V., Alkire, M. B., Ashik, I. M., Baumann, T. M., Carmack, E. C., Goszczko, I., Guthrie, J., Ivanov, V. V., Kanzow, T., Krishfield, R., Kwok, R., Sundfjord, A., Morison, J., Rember, R., & Yulin, A. (2017). Greater role for Atlantic inflows on sea-ice loss in the Eurasian Basin of the Arctic Ocean. Science, 356, 285–291. https://doi.org/10.1126/science.aai8204

75. Post, E., Bhatt, U. S., Bitz, C. M., Brodie, J. F., Fulton, T. L., Hebblewhite, M., Kerby, J., Kutz, S. J., Stirling, I., & Walker, D. A. (2013). Ecological consequences of sea-ice decline. Science, 341, 519–524. https://doi.org/10.1126/science.1235225

76. Poulin, M., Daugbjerg, N., & Gradinger, R. (2011). The pan-Arctic biodiversity of marine pelagic and sea-ice unicellular eukaryotes: a first-attempt assessment, 13–28. https://doi.org/10.1007/s12526-010-0058-8

77. Poulin, M., Underwood, G. J., & Michel, C. (2014). Sub-ice colonial *Melosira arctica* in Arctic first-year ice. Diatom Research, 29, 213–221. https://doi.org/10.1080/0269249X.2013.877085

78. R Core Team. (2021). R: A language and environment for statistical computing. R Foundation for Statistical Computing. Vienna, Austria. https://www.R-project.org/

79. Randelhoff, A., Oziel, L., Massicotte, P., Bécu, G., Lacour, L., Dumont, D., Vladoiu, A., Marec, C., Bruyant, F., Houssais, M.-N., Tremblay, J.-é., Deslongchamps, G., & Babin, M. (2019). The evolution of light and vertical mixing across a phytoplankton ice-edge bloom. Elementa: Science of the Anthropocene, 7, 1–19.

80. Renaut, S., Devred, E., & Babin, M. (2018). Northward Expansion and Intensification of Phytoplankton Growth During the Early Ice-Free Season in Arctic. Geophysical Research Letters, 45, 10, 590–10, 598. https://doi.org/10.1029/2018GL078995

81. Ribeiro, C. G., Lopes, A., Gourvil, P., Gall, F. L., Marie, D., Tragin, M., Probert, I., & Vaulot, D. (2020). Culturable diversity of Arctic phytoplankton during pack ice melting. Elementa: Science of the Anthropocene, 8.

82. Saint-Béat, B., Fath, B. D., Aubry, C., Colombet, J., Dinasquet, J., Fortier, L., Galindo, V., Grondin, P.-L., Joux, F., Lalande, C., LeBlanc, M., Raimbault, P., Sime-Ngando, T., Tremblay, J.-E., Vaulot, D., Maps, F., Babin, M., Deming, J. W., & Bowman, J. (2020). Contrasting pelagic ecosystem functioning in eastern and western Baffin Bay revealed by trophic network modeling. Elementa: Science of the Anthropocene, 8, 1–24. https://doi.org/10.1525/elementa.397

83. Sakshaug, E. (2004). Primary and Secondary Production in the Arctic Seas, In The organic carbon cycle in the arctic ocean. Berlin. https://doi.org/10.1007/978-3-642-18912-8_3

84. Schiffrine, N., Tremblay, J. É., & Babin, M. (2020). Growth and Elemental Stoichiometry of the Ecologically-Relevant Arctic Diatom Chaetoceros gelidus: A Mix of Polar and Temperate. Frontiers in Marine Science, 6, 1–15. https://doi.org/10.3389/fmars.2019.00790

85. Schmidt, K., Brown, T., Belt, S., Ireland, L., Taylor, K. W., Thorpe, S., Ward, P., & Atkinson, A. (2018). Do pelagic grazers benefit from sea ice? Insights from the Antarctic sea ice proxy IPSO25. Biogeosciences, 15, 1987–2006. https://doi.org/10.5194/bg-15-1987-2018

86. Schoemann, V., Becquevort, S., Stefels, J., Rousseau, V., & Lancelot, C. (2005). *PHAEOCYSTIS* BLOOMS IN THE GLOBAL OCEAN AND THEIR CONTROLLING MECHANISMS: A REVIEW. Journal of Sea Research, 53, 43–66. https://doi.org/10.1016/j.seares.2004.01.008

87. Serreze, M. C., Holland, M. M., & Stroeve, J. (2007). Perspectives on the Arctic’s shrinking sea-ice cover. Science, 315, 1533–1536. https://doi.org/10.1126/science.1139426

88. Simon, N., Foulon, E., Grulois, D., Six, C., Desdevises, Y., Latimier, M., Le Gall, F., Tragin, M., Houdan, A., Derelle, E., Jouenne, F., Marie, D., Le Panse, S., Vaulot, D., & Marin, B. (2017). Revision of the genus Micromonas Manton et Parke (Chlorophyta, Mamiellophyceae), of the type species M. pusilla (Butcher) Manton & Parke and of the species M. commoda van Baren, Bachy and Worden and description of two new species based on the genetic and phenotypic characterization of cultured isolates. Protist, 168, 612–635. https://doi.org/10.1016/j.protis.2017.09.002

89. Sjöqvist, C. O., & Kremp, A. (2016). Genetic diversity affects ecological performance and stress response of marine diatom populations. ISME Journal, 10, 2755–2766. https://doi.org/10.1038/ismej.2016.44

90. Smyth, T. J., Tyrell, T., & Tarrant, B. (2004). Time series of coccolithophore activity in the Barents Sea, from twenty years of satellite imagery. Geophysical Research Letters, 31, 2–5. https://doi.org/10.1029/2004GL019735

91. Stroeve, J., Markus, T., Boisvert, L., Milller, J., & Barret, A. (2014). Changes in Arctic melt season and implications for sea ice loss. Geophysical Research Letters, 41, 1216–1225. https://doi.org/10.1002/2013GL058951.Received

92. Tang, C. C., Ross, C. K., Yao, T., Petrie, B., DeTracey, B. M., & Dunlap, E. (2004). The circulation, water masses and sea-ice of Baffin Bay. Progress in Oceanography, 63, 183–228. https://doi.org/10.1016/j.pocean.2004.09.005

93. Tedesco, L., Vichi, M., & Scoccimarro, E. (2019). Sea-ice algal phenology in a warmer Arctic. Science Advances, 5. https://doi.org/10.1126/sciadv.aav4830

94. Tragin, M., & Vaulot, D. (2019). Novel diversity within marine Mamiellophyceae (Chlorophyta) unveiled by metabarcoding. Scientific Reports, 9, 1–14. https://doi.org/10.1038/s41598-019-41680-6

95. Trefault, N., De la Iglesia, R., Moreno-Pino, M., Lopes dos Santos, A., Gérikas Ribeiro, C., Parada-Pozo, G., Cristi, A., Marie, D., & Vaulot, D. (2021). Annual phytoplankton dynamics in coastal waters from Fildes Bay, Western Antarctic Peninsula. Scientific Reports, 11, 1–16. https://doi.org/10.1038/s41598-020-80568-8

96. Vader, A., Marquardt, M., Meshram, A. R., & Gabrielsen, T. M. (2015). Key Arctic phototrophs are widespread in the polar night. Polar Biology, 38, 13–21. https://doi.org/10.1007/s00300-014-1570-2

97. Vaulot, D., Sim, C. W. H., Denise, W., Teo, B., Biwer, C., Jamy, M., & Lopes dos Santos, A. (2022). metaPR 2: a database of eukaryotic 18S rRNA metabarcodes with an emphasis on protists. bioRxiv, preprint, 1–16.

98. Verity, P. G., Brussaard, C. P., Nejstgaard, J. C., Van Leeuwe, M. A., Lancelot, C., & Medlin, L. K. (2007). Current understanding of Phaeocystis ecology and biogeochemistry, and perspectives for future research. Biogeochemistry, 83, 311–330. https://doi.org/10.1007/978-1-4020-6214-8_21

99. Vilgrain, L., Maps, F., Picheral, M., Babin, M., Aubry, C., Irisson, J. O., & Ayata, S. D. (2021). Trait-based approach using in situ copepod images reveals contrasting ecological patterns across an Arctic ice melt zone. Limnology and Oceanography, 66, 1155–1167. https://doi.org/10.1002/lno.11672

100. Wassmann, P., Reigstad, M., Haug, T., Rudels, B., Carroll, M. L., Hop, H., Gabrielsen, G. W., Falk-Petersen, S., Denisenko, S. G., Arashkevich, E., Slagstad, D., & Pavlova, O. (2006). Food webs and carbon flux in the Barents Sea. Progress in Oceanography, 71, 232–287. https://doi.org/10.1016/j.pocean.2006.10.003

101. Wickham, H. (2016). Ggplot2: Elegant graphics for data analysis. https://ggplot2.tidyverse.org

102. Wickham, H., Averick, M., Bryan, J., Chang, W., McGowan, L. D., François, R., Grolemund, G., Hayes, A., Henry, L., Hester, J., Kuhn, M., Pedersen, T. L., Miller, E., Bache, S. M., Müller, K., Ooms, J., Robinson, D., Seidel, D. P., Spinu, V., … Yutani, H. (2019). Welcome to the tidyverse. Journal of Open Source Software, 4, 1686. https://doi.org/10.21105/joss.01686

103. Xu, D., Kong, H., Yang, E.-j., Li, X., Jiao, N., Warren, A., Wang, Y., Lee, Y., Jung, J., & Kang, S.-h. (2020). Contrasting Community Composition of Active Microbial Eukaryotes in Melt Ponds and Sea Water of the Arctic Ocean Revealed by High Throughput Sequencing. Frontiers in Microbiology, 11, 1–15. https://doi.org/10.3389/fmicb.2020.01170

104. Zingone, A., Chrétiennot-Dinet, M. J., Lange, M., & Medlin, L. (1999). Morphological and genetic characterization of *Phaeocystis cordata* and *P. jahnii* (Prymnesiophyceae), two new species from the Mediterranean Sea. Journal of Phycology, 35, 1322–1337. https://doi.org/10.1046/j.1529-8817.1999.3561322.x

105. Zweng, M. M., & Münchow, A. (2006). Warming and freshening of Baffin Bay, 1916-2003. Journal of Geophysical Research: Oceans, 111, 1–13. https://doi.org/10.1029/2005JC003093

